# Inferring cancer type-specific patterns of metastatic spread using Metient

**DOI:** 10.1101/2024.07.09.602790

**Authors:** Divya Koyyalagunta, Karuna Ganesh, Quaid Morris

**Affiliations:** Tri-Institutional Graduate Program in Computational Biology and Medicine, Weill Cornell Medicine, New York, NY 10065, USA; Computational and Systems Biology Program, Sloan Kettering Institute, New York, NY 10065, USA; Department of Medicine, Memorial Sloan Kettering Cancer Center, New York, NY, USA; Molecular Pharmacology Program, Sloan Kettering Institute, Memorial Sloan Kettering Cancer Center, New York, NY, USA

**Keywords:** migration history inference, metastasis, combinatorial optimzation

## Abstract

Cancers differ in how they establish metastases. These differences can be studied by reconstructing the metastatic spread of a cancer from sequencing data of multiple tumors. Current methods to do so are limited by computational scalability and rely on technical assumptions that do not reflect current clinical knowledge. Metient overcomes these limitations using gradient-based, multi-objective optimization to generate multiple hypotheses of metastatic spread and rescores these hypotheses using independent data on genetic distance and organotropism. Unlike current methods, Metient can be used with both clinical sequencing data and barcode-based lineage tracing in preclinical models, enhancing its translatability across systems. In a reanalysis of metastasis in 169 patients and 490 tumors, Metient automatically identifies cancer type-specific trends of metastatic dissemination in melanoma, high-risk neuroblastoma, and non-small cell lung cancer. Its reconstructions often align with expert analyses but frequently reveal more plausible migration histories, including those with more metastasis-to-metastasis seeding and higher polyclonal seeding, offering new avenues for targeting metastatic cells. Metient’s findings challenge existing assumptions about metastatic spread, enhance our understanding of cancer type-specific metastasis, and offer insights that inform future clinical treatment strategies of metastasis.

## Introduction

Metastasis is associated with 90% of cancer deaths, yet its pathophysiology remains poorly understood ^1^. It remains unclear how often clones seed metastases polyclonally, or how often metastases are capable of seeding other metastases, such as through intermediate lymph nodes ^2–10^. It is also not known whether metastatic potential is rare, and thus gained once in the same cancer, or common, and thus gained multiple times ^11–14^. The answers to all these questions would improve the understanding and clinical management of metastasis, and could be gleaned from reconstructing migration histories of metastatic clones from clinical sequencing data.

However, this reconstruction requires solving a complex combinatorial optimization problem ^2–4^, which previous algorithms have addressed by solving a more tractable, oversimplified maximum parsimony problem ^5,15–18^. Early approaches framed migration history reconstruction as a classical phylogenetic problem of labeling interior nodes based on leaf node labels ^19–21^, thereby assuming that the optimal reconstruction minimized the number of migrating clones. While these methods scale well to large datasets, such as those generated by lineage tracing experiments ^22^, they often yield unrealistic reconstructions, including extensive reseeding between metastatic sites ^17^. MACHINA ^17^ replaced the simple parsimony model with multi-objective parsimony criteria that also incorporate the number of seeding sites and polyclonal migration events when scoring histories. MACHINA generates more biologically plausible migration histories on small datasets, but its computational complexity makes its unusable on larger datasets, such as those generated by barcode-based single-cell lineage tracing.

Another key limitation is that existing algorithms only return a single migration history, even when multiple equally parsimonious solutions exist. Using multi-objective parsimony scoring methods can also lead to conflicts where different solutions are most parsimonious under different weightings of the parsimony criteria, and currently these conflicts are resolved using pre-defined, ad hoc criteria ^15–17^. For example, one common assumption is that metastases can only be seeded from the primary tumor ^14^. However, such assumptions are challenged by emerging clinical data suggesting that metastasis-to-metastasis seeding might be more common. Indeed, one prevailing model in oncology, the “sequential progression model” – which posits that lymph node metastases give rise to distant metastases – is the rationale for surgical removal of lymph nodes ^23^ and a recent phylogenetic analysis found that the sequential model applied to a third of patients in a colorectal cohort ^24^. Pre-deciding primary-only seeding dismisses the possibility of metastasis-to-metastasis seeding before looking at the data. These ad hoc rules used to define parsimony also impact other interpretations of metastatic spread, such as the frequency that sites are seeded by multiple tumor cells. While multiple lineage tracing experiments in animal models have detected that a majority of tumors are seeded polyclonally ^25^, human tumor sequencing studies report lower rates of polyclonality ^7^, here we show this discrepancy is partly due to previous modeling assumptions. By pre-biasing their reconstructions, current algorithms undermine a key goal in metastasis research: determining which patterns of metastatic spread are prevalent in different cancer types.

To address these issues and overcome the limitations of previous tools (Supplementary Table 1), we introduce Metient (**met**astasis + grad**ient**). Metient is a principled statistical algorithm that proposes multiple potential hypotheses of metastatic spread in a patient. Metient combines modern stochastic optimization techniques with metastasis priors, i.e., new biologically-grounded, migration history scoring criteria. These priors allow Metient to navigate trade-offs among competing explanations, align predictions with empirical data, and uncover trends specific to cancer subtypes. Together, these advances allow Metient to resolve the current challenges in scalability, parsimony resolution, and biological plausibility in migration history reconstruction.

On realistic simulations, Metient outperforms MACHINA in reconstructing the ground truth migration history. When applied to patient cohorts with breast, skin, ovarian, neuroblastoma, and lung cancers, Metient automatically identifies all plausible expert-assigned migration histories and, in some cases, uncovers more biologically plausible reconstructions, especially when previous analyses on this same data pre-selected a favored seeding pattern. In these cohorts, Metient reveals that polyclonal metastatic seeding occurs far more frequently than previously reported, leading to more robust, experimentally-aligned estimates of clonality. Notably, Metient is the first computational framework capable of scaling to lineage tracing datasets comprising thousands of single cells while avoiding the restrictive assumptions of simplified migration history inference methods.

Metient is free, open-source software that includes easy-to-use visualization tools to compare multiple hypotheses of metastatic dissemination. Metient is accessible at https://github.com/morrislab/metient/.

## Results

### The Metient algorithm

Migration history inference algorithms aim to reconstruct the spread of cancer clones across anatomical sites using molecular sequencing data from primary and metastatic tumors, paired with an unlabeled tree encoding their genetic ancestry (Figure 1a). These methods estimate the proportions of clonal populations at each site (referred to as “witness nodes”, Figure 1b,c), or use observed cell locations directly (e.g., from lineage tracing data). The interior nodes of the tree are then labeled with anatomical sites, defining migration events — tree edges connecting clones assigned to different sites are deemed “migration edges”. The reconstructed history is referred to as a “migration history” (Figure 1c).

**Figure 1.**
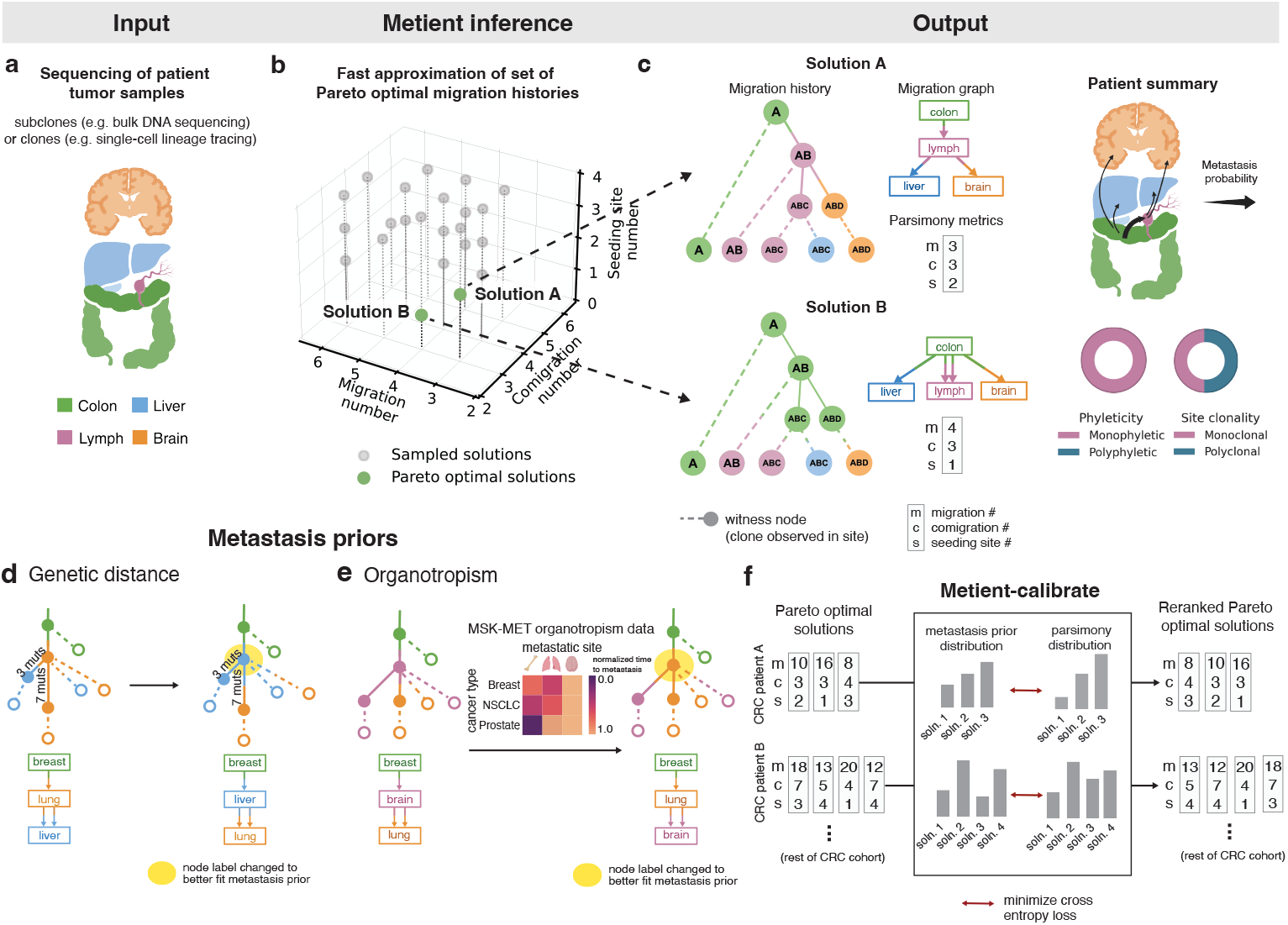
A scalable framework for migration history inference evaluates multiple hypotheses of metastatic spread. (a) Data on clone or subclone frequencies from multiple tissue samples is input into Metient. (b) Metient efficiently recovers multiple Pareto-optimal migration histories based on the three counts: migrations, comigrations and seeding site number. (c) Metient solutions can be represented by a migration history, a tree with (1) an anatomical site labeling of its internal nodes, and (2) leaf nodes (witness nodes) representing a node’s presence in an anatomical site. A migration graph summarizes the migration edges of the migration history. Parsimony metrics indicate the number of migrations (*m*), comigrations (*c*), and seeding sites (*s*). Metient can report summary metrics over the Pareto front, such as the probability of a particular tissue-to-tissue transition, and the percentage of phyleticity and clonality that is inferred. (d) An example of how using genetic distance (a metastasis prior) can promote migration histories with migrations on longer edges with more mutations. The assigned anatomical site label of the highlighted node changes. (e) An example using organotropism (a metastasis prior) to identify migration histories with unlikely metastatic events, such as subsequent metastasis from the brain. The anatomical site label of the highlighted node is changed. (f) Metient-calibrate uses the metastasis priors to fit weights for the parsimony metrics, this calibrated parsimony model can then be used to rescore the Pareto front of migration histories produced for each patient in that cohort.

The most advanced existing algorithm for migration history inference, MACHINA ^17^, scores migration histories using three parsimony metrics: (1) migrations— the number of times a clone is established at a new site ^4,15–17^; (2) comigrations—the number of migration events involving multiple clones traveling together; and (3) seeding sites—the number of sites from which clones migrate. MACHINA searches for the most parsimonious history by minimizing these metrics using mixed-integer linear programming (MILP) ^26^.

While effective, this approach has critical limitations. First, MILP solvers require hard constraints and linear objectives, preventing the integration of complex, biologically-motivated scoring criteria. Second, they identify only a single optimal solution, even when multiple equally plausible histories exist, leading to biases in reconstructions. Finally, MILP methods scale poorly, making them unsuitable for analyzing large trees, such as those arising from single-cell lineage tracing experiments.

Metient addresses these issues with two key innovations. First, it replaces the MILP framework with stochastic optimization, leveraging low-variance gradient estimators to efficiently search the space of possible migration histories (Supplementary Figure S1, Methods, Supplementary Information). Second, it introduces biologically-informed “metastasis priors” to resolve ambiguities in equally parsimonious solutions and resolve trade-offs among migrations, comigrations, and seeding sites (Figure 1c-e) in scoring parsimony. The first step in Metient’s inference is to define a Pareto front ^27^ of potential solutions by searching for parsimonious migration histories under a wide range of “parsimony models” (Supplementary Table 2) represented by a set of weights – *w*_*m*_, *w*_*c*_, and *w*_*s*_ – assigned, respectively, to the number of migrations (indicated by *m*), comigrations (*c*), and seeding sites (*s*). A migration history’s parsimony score, *p*, is the model-weighted average of these three parsimony metrics, i.e., *p* = *w*_*m*_*m* + *w*_*c*_*c* + *w*_*s*_*s*. Different parsimony models favor different histories on the Pareto front.

### Metient-calibrate fits cancer type-specific parsimony models

To assess the importance of considering multiple hypotheses of metastatic spread, we defined four different cancer type-specific patient cohorts consisting of genomic sequencing of matched primary and multiple metastases: melanoma ^3^, high-grade serous ovarian cancer (HGSOC) ^4^, high-risk neuroblastoma (HR-NB) ^9^, and non-small cell lung cancer (NSCLC) ^14^. After quality control (Supplementary Information), our dataset included 479 tumors (143 with multi-region sampling) from 167 patients (melanoma: n=7, HGSOC: n=7, HR-NB: n=27, NSCLC: n=126). Among patients with multiple metastases, 38.2% of patients (29/76) had multiple Pareto-optimal migration histories; this frequency increases to 53% for patients with three or more metastases. When it returns a single solution, Metient has ruled out all other sampled solutions as suboptimal, enhancing confidence in the single Pareto-optimal history. Different Pareto-optimal histories often represented different overall patterns of metastatic spread. For example, Figure 1c shows a patient with metastatic colorectal cancer with two Pareto-optimal reconstructions: one in which a lymph node metastasis gives rise to all other metastatic tumors (solution A), and another where most metastases are seeded directly from the primary tumor (solution B). Here, forcing an arbitrary choice between the two reconstructions determines whether one concludes that the lymph node acted as a staging site for metastatic spread.

MACHINA resolves parsimony conflicts by prioritizing the minimization of migrations, comigrations, and seeding sites in a fixed hierarchical manner (i.e., *w*_*m*_ >> *w*_*c*_ >> *w*_*s*_). Other methods simplify further, focusing solely on minimizing migrations ^4,15,22^. Such a rigid model can obscure important differences among cancers. For example, in ovarian cancer metastatic events are often polyclonal due to the dissemination of clusters of metastatic cells through peritoneal fluid, these trends would lead to more migrations relative to comigrations ^28–30^. In contrast, multiple seeding sites might be more common in the estimated 23.4% of cancer cases have lymph node involvement ^31^, particularly in cancer types like non-small cell lung cancer ^32^ where this frequently increases up to 87%. It has been proposed that metastatic cells can make a “pit stop” at regional lymph nodes before disseminating to other distant sites ^33^. Prescribing a single, cancer-independent parsimony model risks missing these clinically-relevant differences.

In contrast to an ad hoc approach, Metient uses metastasis priors to both define a cancer type-specific parsimony model and to resolve ties between histories with equal parsimony metrics. Metient, in calibrate mode, fits a patient cohort-specific parsimony model using the metastasis priors (Figure 1d-f; Methods). This calibrated model is used to rank Pareto-optimal histories that differ in their parsimony metrics. Metient also provides a pan-cancer parsimony model, calibrated to all four cohorts combined, for use when an appropriate patient cohort is not available. This pan-cancer model is a first step toward estimating the likelihoods of key events in metastatic dissemination across cancers, providing a generalized framework that can be applied to any dataset.

Metient provides two metastasis priors. One, genetic distance, can be applied to any cohort. The other, organotropism, can be used when appropriate tissue-type information are available for the sequenced tumor samples. The genetic distance prior considers the genetic distance of migration edges in the labeled clone tree; here genetic distance is defined as the number of mutations gained in the child clone and not present in the parent clone. This prior favors histories with higher averaged genetic distances on migration edges (Figure 1d, Methods, Supplementary Information). Theoretical and empirical evidence supports this approach. Colonizing clones at metastatic sites typically undergo clonal expansions ^34^, making their mutations more detectable than those in the source tumor, where private mutations often remain undetected due to insufficient clonal expansion. Furthermore, metastasizing cells face strong selection pressures, which are associated with elevated mutation rates ^35–37^. These pressures, combined with the increased genomic instability observed in metastases ^35–37^, result in higher tumor mutation burdens compared to primary tumors ^31,38,39^. Analysis of our cohorts confirm this trend, with migration edges showing higher genetic distances than non-migration edges when the most parsimonious migration history was unambiguous (Supplementary Figure S2).

The second metastasis prior, organotropism, is derived from data from over 25,000 metastatic cancer patients to compute the observed preference of certain cancer types for specific metastatic sites ^40^. All else being equal, this prior favors more common migration patterns, like migration from a breast-seeded lung metastasis to the brain, over rarer ones, like a breast-seeded brain metastasis to the lung ^41^ (Figure 1e). The organotropism prior does not directly model secondary metastasis-to-metastasis migrations, instead it indirectly captures these patterns through its scoring of primary-to-metastasis events (Methods).

Metient uses these priors to complement, rather than replace, the parsimony model. In our benchmarking analyses on simulated data, we find that using genetic distance alone to score migration histories performs poorly and can result in the inference of highly non-parsimonious migration histories (Supplementary Tables 3, 4, see also PathFinder ^18^). Therefore, genetic distance and organotropism are used to rank solutions only after defining the Pareto front. Metient reports and visualizes all Pareto-optimal solutions it finds, allowing users to evaluate multiple plausible hypotheses and select the one most consistent with biological and clinical insights.

### Simulated data validates the genetic distance prior and shows that Metient is state-of-the-art

To assess Metient’s new objective and gradient-based optimization on data with a provided ground-truth, we ran benchmarking analyses along with the state-of-the-art migration history inference method (MACHINA ^17^) on simulated data for 80 patients with 5-11 tumor sites and various patterns of metastatic spread. These data were originally used to validate MACHINA.

First, to assess the added value of the genetic distance prior, we used Metient-calibrate to fit a calibrated parsimony model, and compared calibrated Metient with a version of Metient that used the parsimony model implied by MACHINA. We fit two calibrated models, one on a cohort with primary-only seeding and another on a cohort with metastasis-to-metastasis seeding. Metient-calibrate improved recovery of the ground truth migration graph (Supplementary Figure S3a) over the fixed parsimony model (Calibrate vs. Evaluate (MP) in Supplementary Table 3), showcasing the ability of the metastasis priors to learn metastatic patterns specific to a cohort and improve overall accuracy. In addition, Metient-calibrate predicts ground truth seeding clones and migrations graphs at least as accurately as MACHINA, with overall improvements as tree sizes get larger (Supplementary Figure S3b) and significant improvements in inferring the seeding clones for patients with more complex metastasis-to-metastasis seeding (Supplementary Figure S3b right; p=0.0021).

Notably, even though the Metient framework is non-deterministic, it identifies the same top solution 97% of the time across multiple runs (Supplementary Figure S3c). In addition to its improved accuracy, Metient runs up to 55x faster (3.95s with Metient-64 vs. 221.19s with MACHINA for a cancer tree with 18 clones and 9 tumors) on this benchmark, showcasing our framework’s scalability as tree sizes grow (Supplementary Figure S3d).

### Multi-cancer analysis of clonality, phyleticity, and dissemination patterns

After validating calibrated Metient’s ability to recover ground-truth metastatic patterns in simulated data, we applied it to the patient cohorts with melanoma, HGSOC, HR-NB, and NSCLC to explore shared and unique patterns of metastatic spread. Due to missing or inadequate anatomical site labels, we only calibrated to genetic distance.

We examined three aspects of metastatic dissemination: seeding pattern (single-source from the primary or another site, multi-source, reseeding; Figure 2a), clonality (number of clones seeding metastases; Figure 2b), and phyleticity (metastatic potential is gained in one or multiple evolutionary trajectories; Figure 2c; Methods). We distinguish between genetic polyclonality (multiple clones seeding metastases) and site polyclonality (multiple clones seeding individual sites), to highlight cases where each metastasis is seeded by a single clone, but different sites may be seeded by different clones, in order to distinguish cases where different site-specific mutations are needed for metastasis (Figure 2b).

**Figure 2.**
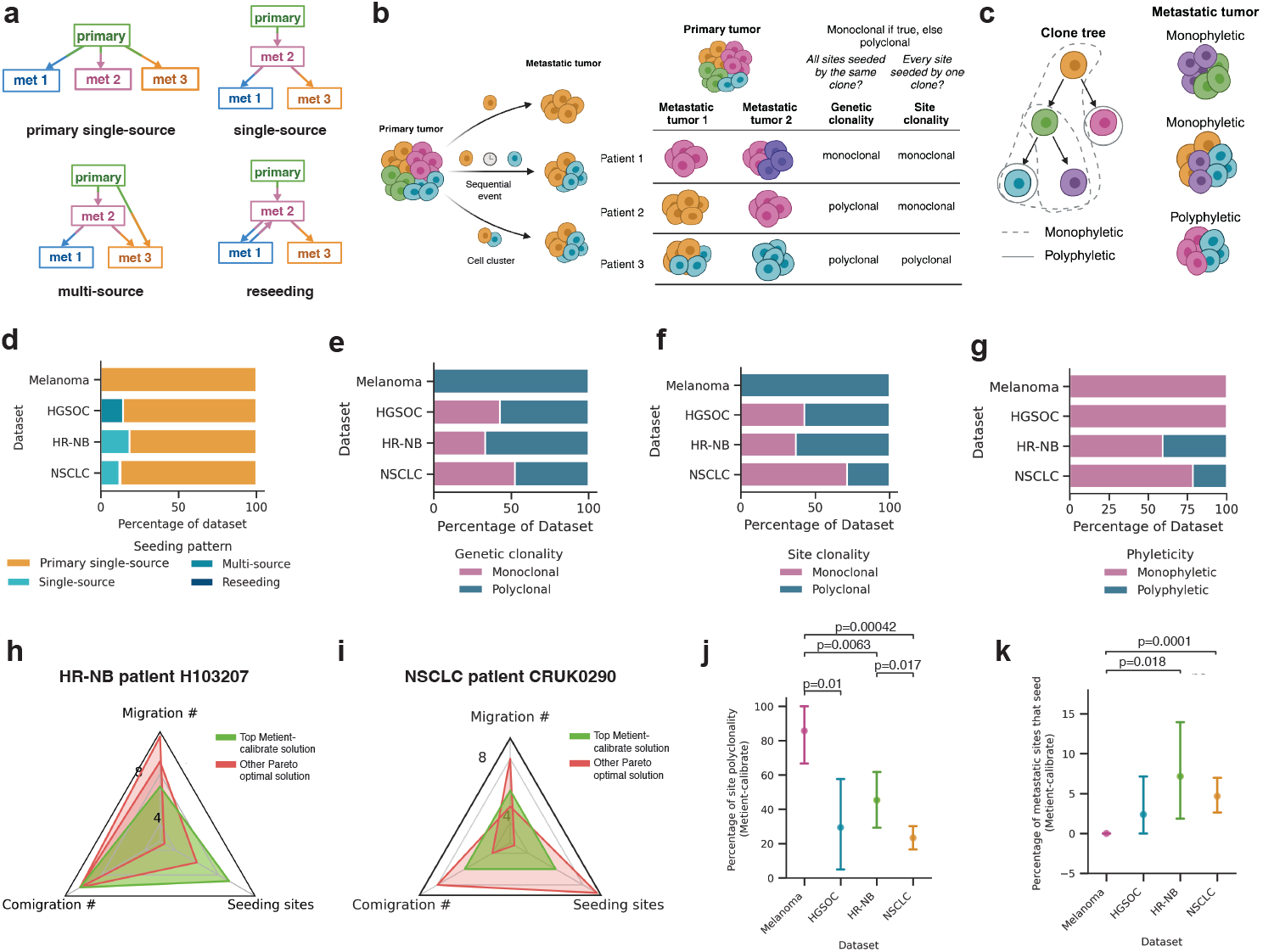
Multi-cancer analysis of clonality, phyleticity, and dissemination patterns. **(a)** Schematic illustrates four metastatic seeding patterns. met: metastasis. **(b)** Schematic illustrates metastases seeding by one or multiple clones either sequentially or in a cell cluster, table defines genetic clonality versus site clonality. Colors represent genetically distinct cancer cell populations. **(c)** Schematic illustrates the definitions of monophyletic and polyphletic seeding. Monophyletic indicates that the colonizing clone closest to the root can reach every other colonizing clone on the clone tree. Colors represent genetically distinct cancer cell populations. Distribution of **(d)** seeding patterns, **(e)** genetic clonality, **(f)** site clonality and **(g)** phyleticity for each dataset, as inferred by Metient’s top migration history. **(h)** Radar plot showing the unique Pareto-optimal metrics for migration histories inferred by Metient for HR-NB patient H103207. **(i)** Radar plot showing the unique Pareto-optimal metrics for migration histories inferred by Metient for NSCLC patient CRUK290. **(j)** Comparing across datasets the percent of migrations that are polyclonal for the top Metient solution. Statistical significance assessed by a Welch’s t-test. Error bars are the standard error for each dataset. **(k)** Comparing across datasets the percent of metastatic sites that seed for the top Metient solution. Statistical significance assessed by a Welch’s t-test. Error bars are the standard error for each dataset.

Consistent with expert annotations ^3,4,9,14,17^, Metient finds that single-source seeding from the primary tumor is the most common pattern in every cohort (Figure 2d). However, Metient identifies a larger fraction of polyclonal migration patterns than previous reports ^8,14^: 53.3% of patients show genetic polyclonality (Figure 2e), and 38.3% of patients have site polyclonality (Figure 2f). Overall, Metient estimates that 34.1% of sites (107/314) are seeded by multiple clones; nearly double previous estimates of site polyclonality (19.2%) based on an analysis of breast, colorectal and lung cancer patients ^8^. Notably, the choice of parsimony model affects polyclonality, as reducing the number of seeding sites increases polyclonal migrations (Supplementary Figure S4a). However, Metient’s findings do not stem from assuming primary-only seeding, as done in prior work.

Metient’s phyleticity estimates are similar to previous reports: 77.2% of patients (129/167) have a monophyletic tree where metastatic potential is gained once and maintained (Figure 2g). For some patients, this is due to the root clone being observed in one or more metastatic sites (Supplementary Figure S4b), and for other patients, all colonizing clones belong to a single subtree of the clone tree whose root is itself a colonizing clone. Either scenario suggests that metastatic potential is less likely to be gained via multiple, independent evolutionary trajectories across cancers.

### Cancer type-specific metastasis trends

We next examined cancer type-specific differences in metastatic trends, first using a bootstrapping approach to ensure that the parsimony metric weights were reproducible and reflective of population level patterns for a particular cancer type. We fit parsimony metric weights to 100 bootstrapped samples of patients within the cohort (Methods), and found that 98.4% of patients ranked the same top solution across bootstrap samples, indicating that Metient can learn a reproducible cancer type-specific model for the melanoma and HGSOC cohorts which have only seven patients each.

These cancer type-specific parsimony metric weights lead to cohort-specific choices on how Metient ranks a patient’s Pareto front of migration histories. For example, Metient chooses the solution on the Pareto front with lowest migration number (i.e. colonizing clones) for HR-NB patient H103207 (Figure 2h), but the solution with the median value of each metric for NSCLC patient CRUK0290 (Figure 2i). To systematically assess the impact of cohort-specific rankings we computed the percentage of polyclonality and number of seeding sites in the top ranked solution for patients with each cancer type. Overall, we found a significantly higher fraction of polyclonal migrations in melanoma than HGSOC, HR-NB and NSCLC patients (Figure 2j). One explanation for this heightened polyclonality in melanoma patients is that all patients in the cohort had locoregional skin metastases, a common “in-transit” metastatic site around the primary melanoma or between the primary melanoma and regional lymph nodes. These locoregional sites could have multiple cancer cells traveling together through hematogeneous or lymphatic routes to seed new localized tumors ^42^. The HR-NB and NSCLC cohorts had significantly higher percentages of metastasis-to-metastasis seeding than melanoma (Figure 2k). As described below, in the HR-NB cohort, multiple patients exhibit metastasis-to-metastasis seeding within an organ or between commonly metastatic sites. In the NSCLC cohort, 76.2% of patients have lymph node metastases, from which it is known that further metastases are commonly seeded ^43^. Indeed, Metient predicted that 75% (12/16) of NSCLC patients with metastasis-to-metastasis seeding had seeding from a lymph node to other metastases, suggesting that lymph nodes serve as a common intermediate staging site.

### Metastasis priors identify biologically relevant migration histories and alternative explanations of spread

A core advance of Metient is its ability to identify and rank the Pareto-optimal histories of a patient’s cancer. To assess how well our top ranked solution aligns with the most biologically plausible explanation, we compared our inferred migration histories to previously reported, expert-annotated seeding patterns. Of the 167 patients analyzed, 152 patients had an expert or model-derived annotation available. In 84% of cases (128/152), Metient’s predictions aligned with previously reported site-to-site migrations. For the remaining 24 cases, Metient either identified a more parsimonious history or included the expert annotation on the Pareto front but prioritized a different solution based on its metastasis priors. We provide a detailed case-by-case comparison in the Supplementary Information and Supplementary Figures S5, S6, S7, S8, and highlight two interesting HR-NB cases below.

Metient predicts metastasis-to-metastasis seeding for two HR-NB cases (H103207, H132384), differing from earlier reports of direct seeding from the primary ^9^. HR-NB patient H103207 exhibits evidence for two potential metastasis-to-metastasis seeding scenarios, both involving seeding between the lung and liver (Figure 3a). While the exact prevalence of metastasis-to-metastasis seeding between the liver and lung in HR-NB is unknown, both are common sites of metastases across cancer types due to cancer cells’ ability to take advantage of rich blood supply and vascular organization ^40^. Colonization of the lung by clones from a primary liver tumor is common ^40,44,45^ and, similarly, the liver is a common site of metastasis for primary lung cancer patients ^40,46^, suggesting that transitions from a liver-competent cancer clone to a lung-competent one or vice versa could be common as well. The lung- and liver-colonizing clones emerge on a shared branch, separate from the branch that gives rise to the CNS-colonizing clones (Supplementary Figure S5a). This suggests that evolution within the primary tumor generated clones with shared metastatic potential for the lung and liver that was separate from the metastatic potential of CNS-colonizing clones. This is consistent with the clonal analysis reported by Gundem et al. ^9^, and further supports the proposed metastasis-to-metastasis seeding. Patient H132384 also shows evidence of metastasis-to-metastasis seeding, but from bone-to-bone, first to the left cervical and secondarily to the chest wall (Figure 3b). Metastasizing cells exhibit organ-specific genetic and phenotypic changes to survive in a new microenvironment ^40^, suggesting that seeding an additional tumor within the same organ microenvironment is more likely than a secondary migration from the primary adrenal tumor in this case. In addition, prior experimental evidence shows that bone metastases prime and reprogram cells to form further secondary metastases ^47,48^. These posited metastasis-to-metastasis seedings are thus supported by site proximity or organotropism, or both, and these Metient reconstructions were made without providing such information.

**Figure 3.**
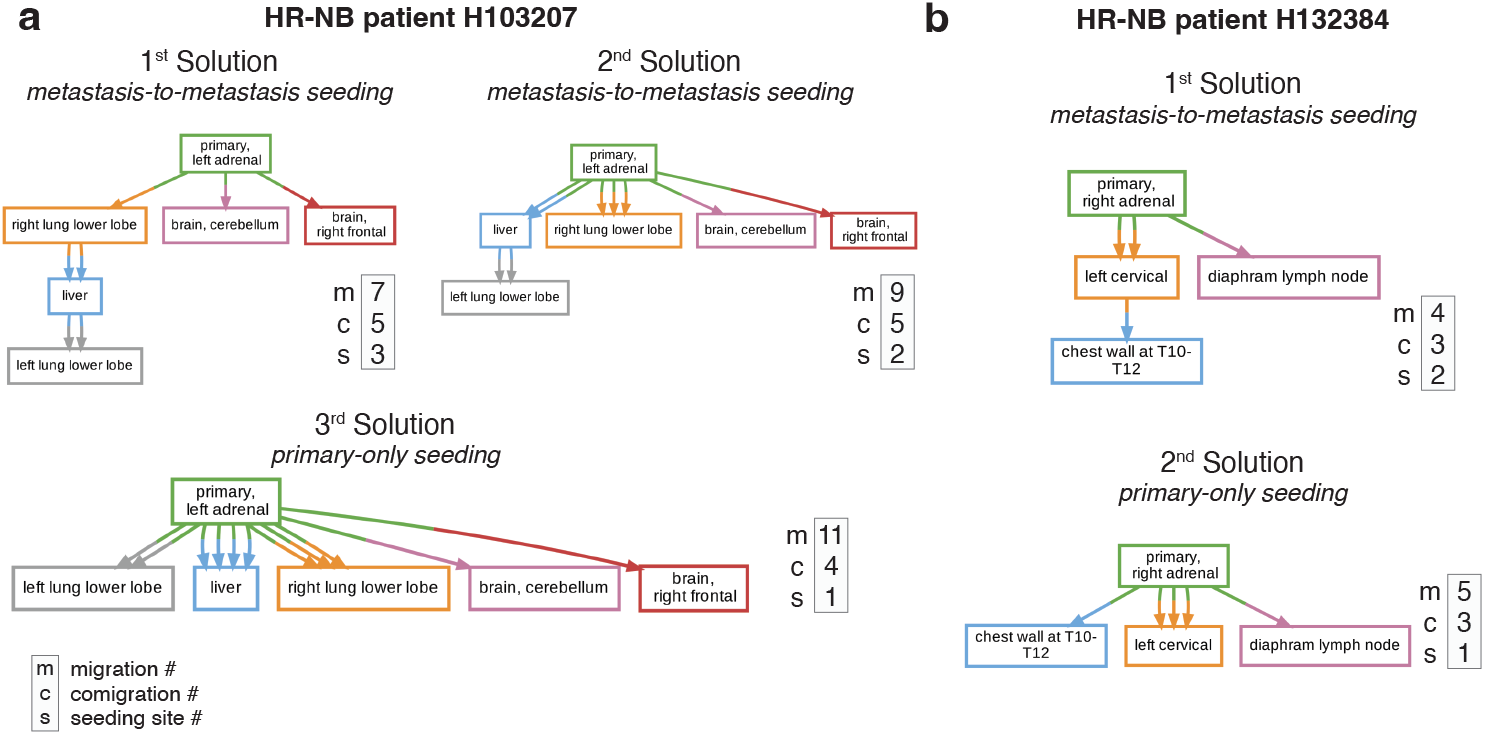
Metient finds biologically relevant migration histories. **(a)** All ranked Pareto-optimal migration graphs inferred by Metient-calibrate for HR-NB patient H103207. **(b)** All ranked Pareto-optimal migration graphs inferred by Metient-calibrate for HR-NB patient H132384.

Next, we compared the inferred migration histories from the NSCLC samples we analyzed to an in-depth analysis of the same samples by the TRACERx consortium ^14^ which enforces a primary single-source dissemination model for its analysis of clonality and phyleticity. While Metient generally agrees with this dissemination model, Metient predicts metastasis-to-metastasis seeding for several (12.8%; 16/126) patients. For instance, Metient suggests that, in patient CRUK0484, a rib metastasis seeded the scapula (Figure 4a), a pattern consistent with other bone-to-bone metastasis cases like HR-NB patient H132384.

**Figure 4.**
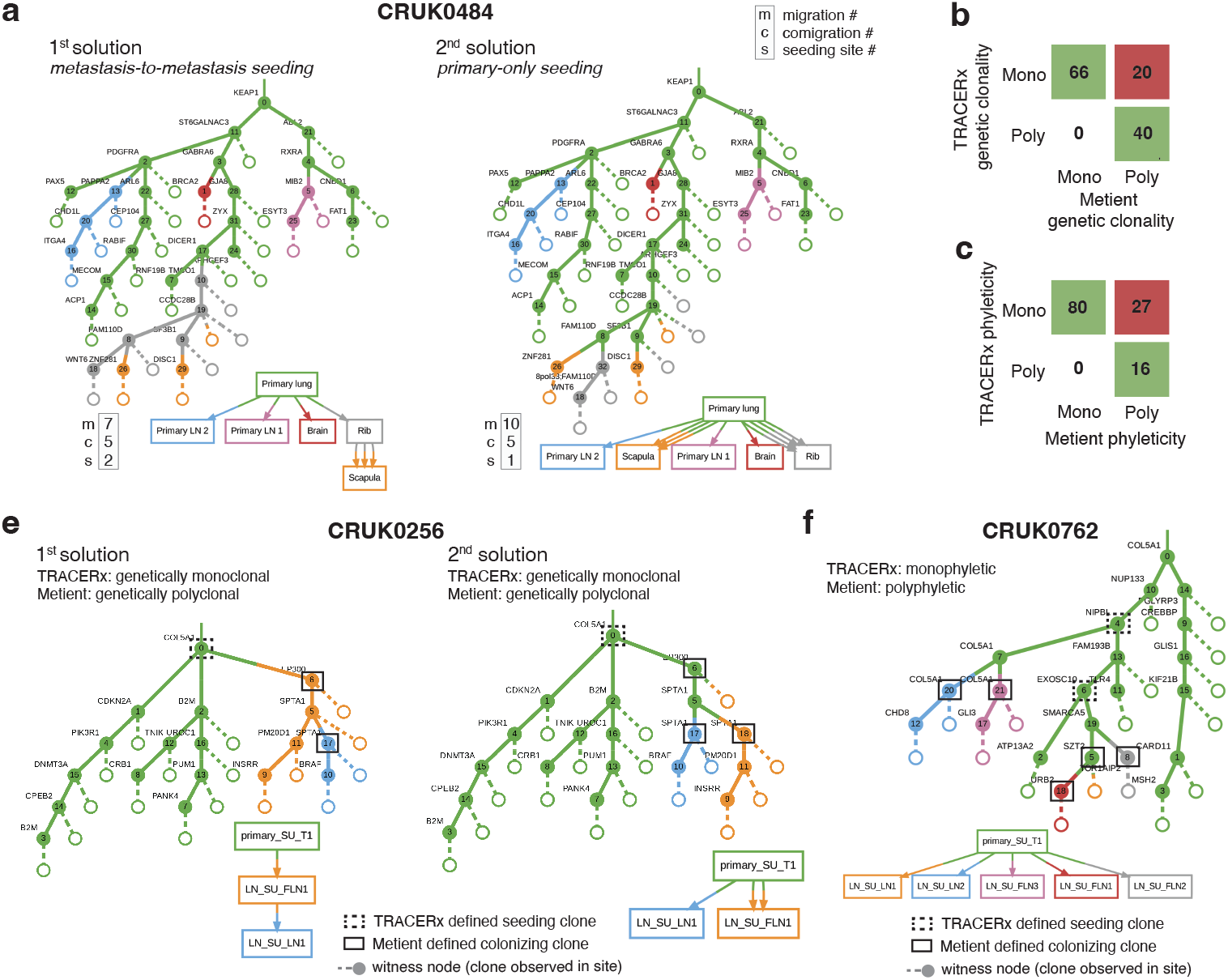
Metient identifies more polyclonality and polyphyelticity in a large-scale NSCLC cohort. **(a)** The top two Pareto-optimal solutions for NSCLC patient CRUK0484 as ranked by Metient-calibrate. Comparison of Metient’s inference to TRACERx’s: **(b)** clonality and **(c)** phyleticity classification. Counts indicate the number of patients in agreement or disagreement. **(d)** All Pareto-optimal solutions for NSCLC patient CRUK0762 as ranked by Metient-calibrate. **(f)** Patient CRUK0762 where seeding pattern and clonality are in agreement between TRACERx and Metient-calibrate but phyleticity differs due to which clones are classified as seeding.

Metient’s highest-scoring solution agrees with TRACERx classifications 84.1% (106/126) for clonality (Figure 4b)) and 78% (96/123) for phyleticity (Figure 4c). Discrepancies arise because TRACERx defines seeding clones as the most recent shared clone between primary and metastasis, while Metient considers the full migration history and accounts for metastasis-to-metastasis seeding, rather than assuming primary-only seeding. Consequently, Metient has higher sensitivity for detecting colonizing clones, leading to increased detection of polyclonal and polyphyletic events.

In 20 NSCLC patients, Metient inferred that multiple colonizing clones are needed to explain the full migration history, whereas no history is consistent with the TRACERx identified colonizing clones. For example, in CRUK0256 (Figure 4d), Metient identifies multiple colonizing clones, in either the metastasis-to-metastasis seeding scenario (Figure 4d first solution) or a primary-only seeding scenario (Figure 4d second solution), whereas TRACERx identifies only the root clone. Similarly, TRACERx’s phyleticity inferences, constrained by its definition of colonizing clones, miss some potential polyphyletic events, misclassifying them as monophyletic. For instance, Metient classifies 27 cases as polyphyletic where TRACERx classifies them as monophyletic, as in patient CRUK0762 (Figure 4e). While monophyleticity remains the dominant pattern in NSCLC (65%), we suggest that polyphyleticity may be underrecognized due to TRACERx’s lower sensitivity for detecting colonizing clones.

### Metient identifies the mediastinum as an early hub for metastatic dissemination in lung adenocarcinoma single-cell lineage tracing experiments

Recent advancements in Cas9-based molecular recording technologies have enabled precise mapping of cellular evolution and metastatic progression in cancer, offering invaluable insights for metastasis research. However, existing multi-objective models fail to scale to the size of this data, which often profiles thousands of single cells, while simplified approaches frequently yield erroneous results. We demonstrate the application of Metient to lineage tracing data from a human *KRAS*-mutant lung adenocarcinoma cell line, tracking 100 clones in a mouse xenograft model ^22^. Metient is the first method capable of analyzing such large datasets—encompassing clones with up to 15,488 cells—without relying on oversimplified models of metastasis.

Using Metient’s pan-cancer parsimony model, calibrated on bulk-sequencing data, we identified diverse patterns on the Pareto front, providing insights into the different possible trajectories of metastatic spread for each clone. For example, clone 15’s Pareto front consisted of 21 unique solutions (Figure 5a). The lowest-ranked solution (Figure 5b) exhibits primary-only seeeding, whereas higher-ranked solutions revealed greater involvement of a mediastinum seeding site, thereby reducing the overall number of migration events (Figure 5c,d). When multiple solutions on the Pareto front had equal parsimony metrics, Metient used the organotropism prior to distinguish between them (see Supplementary Information for details).

**Figure 5.**
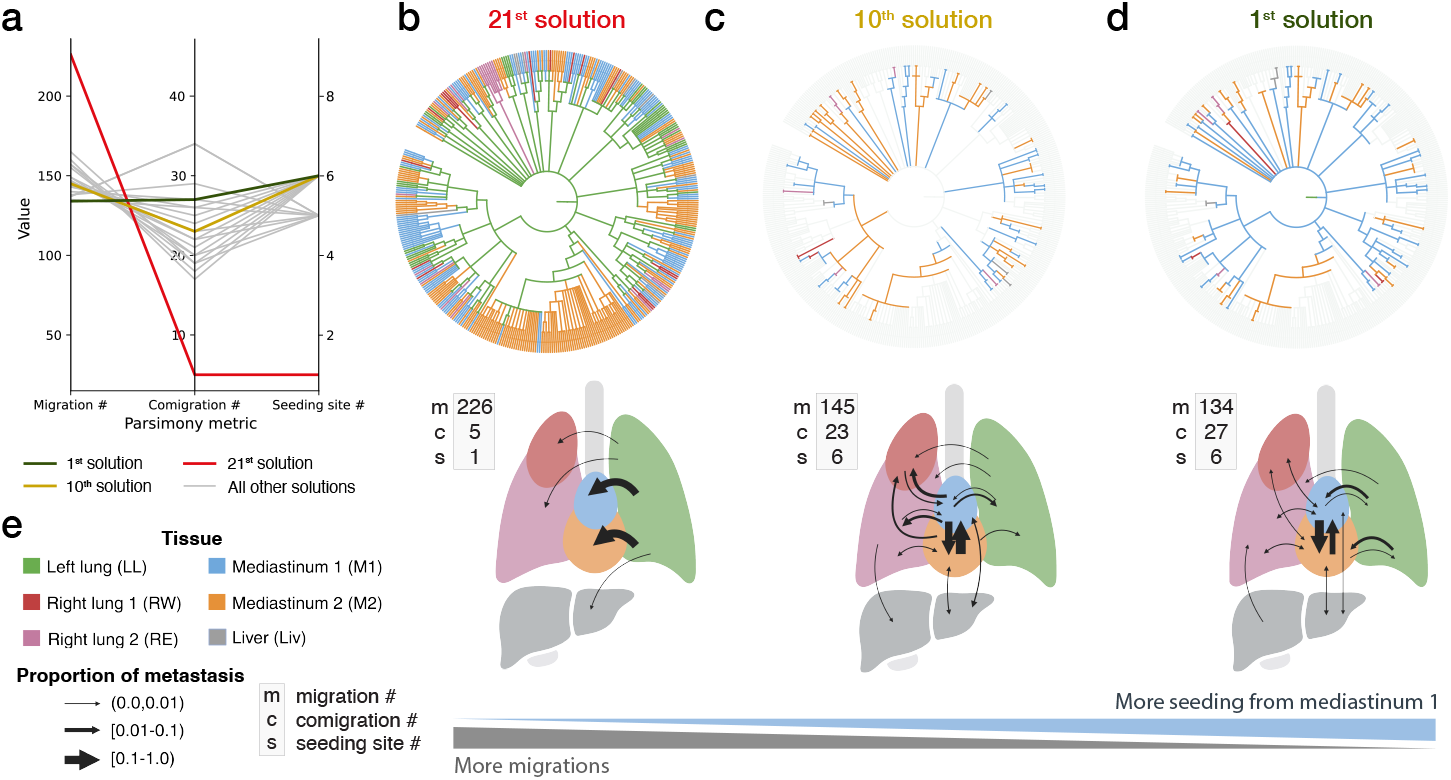
Clones from lineage tracing data have large Pareto fronts with many different possible patterns of spread. (a) The parsimony metrics for all 21 solutions on the Pareto front for clone 15. (b-d) Solutions by their rank on the Pareto front, where more top ranked solutions exhibit lower migration numbers and more seeding via the mediastinum 1. Solutions ranked 1 and 10 only show colored migration edges that are different from the primary-only seeding solution ranked 21. Migration edges that are the same as those in solution ranked 21 are in light grey. (e) Legend for colors and width of arrows used in b-d.

We compared Metient’s migration history inferences for the 100 clones to those inferred by FitchCount ^22^, the method used in the original paper. FitchCount, an extension of the single-objective Fitch-Hartigan algorithm ^20,21^, computes tissue transition probabilities for minimum migration number solutions. However, FitchCount neither recovers complete migration histories nor integrates the more biologically relevant multi-objective parsimony model. Its simplified assumptions led to notable differences in the overall tissue transition probabilities. Specifically, Metient postulates that seeding from the primary left lung tumor predominantly spreads to the mediastinal tissues, which function as a hub for further metastatic dissemination (Figure 6a,b).

**Figure 6.**
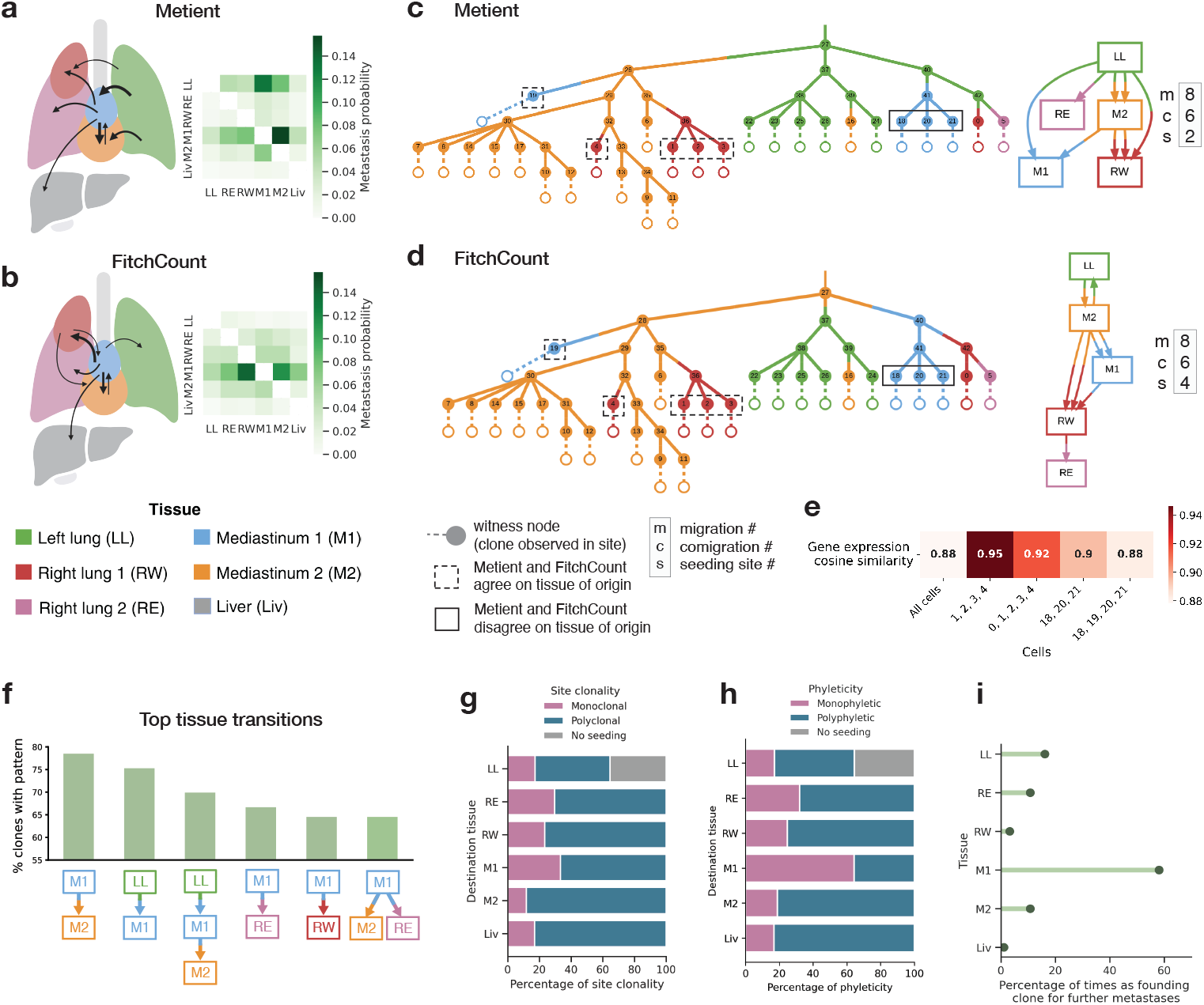
Metient’s migration histories of lineage tracing clones differ from simplified models and implicate the mediastinum as an early metastatic hub. Probability of metastasis from source to destination tissue, aggregated over all 100 clones, as inferred by Metient in (a) and FitchCount in (b). Width of arrows on the body maps are scaled to the frequency of metastasis between sites, as shown in the heat map. Tissue transition probabilities < 0.05 are omitted. (c) Clone 99’s top ranked migration history by Metient. (d) A random minimum migration solution returned by the Fitch-Hartigan algorithm used by FitchCount for clone 99. (e) The mean cosine similarity of gene expression of cells in clone 99. (f) The top six most frequent subgraphs of migration graphs among the 100 clones. (g) The site clonality of seeding to each destination tissue across all 100 clones. (h) The phyleticity of seeding to each destination tissue across all 100 clones. (i) The percentage of times the first colonizing clone is detected in each tissue.

In addition, there are some notable differences in Metient’s and FitchCount’s migration histories. For example, both methods predict a solution with the same number of metastatic migrations for clone 99, but failing to account for comigrations and seeding sites leads FitchCount to propose a more complex reseeding pattern (Figure 6c,d). This example makes clear that minimum migration solutions are not always the simplest or most biologically relevant explanations of the data. To further evaluate the validity of Metient’s migration histories, particularly where Metient and FitchCount diverged, we used the single-cell RNA-sequencing readouts from the lineage tracing data. Since cells alter their gene expression to adapt to new tissue environments ^40,49^, we hypothesized that a cell’s gene expression profile should be more similar to cells that underwent the same metastatic transition—i.e., those originating from the same source tissue and metastasizing to the same final tissue—than to other cells within the same clone. Both Metient’s and FitchCount’s inferred migration histories have significantly increased gene expression cosine similarity for cells that underwent the same metastatic transition (Supplementary Figure S9), with Metient having a slightly higher effect size when the tissue of origin differed from FitchCount (Cohen’s *d* for paired samples: 0.54 for Metient and 0.52 for FitchCount). For example, in clone 99, both methods predict that cells 1-4 were seeded from M2 to RW (Figure 6c,d). As expected, these cells show high gene expression similarity to one another, while the inclusion of cell 0, also in RW but likely originating from a different source, reduces this similarity (Figure 6e). In the same clone, the average gene expression similarity of the group of co-migrating clones (18,20,21) decreases when 19 is added to the group, suggesting that 19 has a different source tissue than clones (18,20,21), as predicted by Metient but not FitchCount (Figure 6c-e).

Metient reconstructs complete migration trees for all Pareto-optimal solutions it finds, allowing for a detailed analysis of migration event sequencing, clonality, and phyleticity. This analysis cannot be done with FitchCount, because it does not return all migration histories it finds, but rather uses a dynamic programming approach that returns one minimum migration number solution along with aggregated counts of tissue transitions over all solutions. In Metient’s reconstructions, mediastinum 1 is the most common source of further dissemination, serving as a hub for systemic spread more often than the primary tumor (Figure 6f). This is consistent with the mediastinal lymph tissue’s anatomical and lymphatic connectivity to the primary lung tumor, as it provides a direct route for cancer cells to metastasize early ^50,51^. The original study also detects early colonization of mediastinal lymph tissue using bulk live imaging ^22^.

Metient revealed widespread polyclonality and polyphyleticity across this dataset, in line with the aggressive metastatic nature of this cell line ^22^ (Figure 6g,h). In a majority of clones, the mediastinum 1 contains the founding clone for further metastases (Figure 6i). This suggests that while subsequent systemic spread is highly polyclonal, the initial seeding of the mediastinum may be driven by a single dominant clone for many cases, suggesting a pivotal role in the early stages of metastatic progression.

## Discussion

We have presented and validated Metient, a novel framework for reconstructing the migration histories of metastasis. In contrast to prior work, Metient defines a Pareto front of possible migration histories, and then uses biologically-motivated metastasis priors to resolve parsimony conflicts in a data-dependent manner. Metient is able to scale to large problems because it adapts Gumbel straight-through stochastic gradient estimation to optimize the combinatorial problem required for history reconstruction. Collectively, these advances improve performance on simulated data, improve biological interpretation on real data, and can define a Pareto front in a fraction of the time that MACHINA, the current state-of-the-art, takes to output a single solution.

We demonstrate that previous algorithms using pre-specified parsimony models and ad hoc rules introduced systematic biases into studies of metastatic spread. For example, a prior analysis of a large NSCLC cohort ^14^ assumed primary-only seeding when assessing clonality and phyleticity. This approach excluded plausible histories involving metastasis-to-metastasis seeding, even when MACHINA was used to identify such events. In contrast, Metient reveals that 12.8% of the NSCLC cohort exhibits evidence of metastasis-to-metastasis seeding, predominantly through lymph nodes. By identifying these overlooked metastatic events and using the full migration history to identify colonizing clones, Metient also uncovers higher levels of polyclonality and polyphyleticity in this cohort than previously reported. Furthermore, across all datasets analyzed, Metient consistently reveals almost twice the level of polyclonality previously reported, suggesting that multiple clones more frequently contribute to metastatic progression. This view of metastatic dynamics implies that therapeutic strategies could shift focus from targeting individual cancer cells to disrupting the formation and migration of tumor cell clusters, potentially offering more effective approaches to prevent or reduce metastasis ^52^. Metient’s statistical, data-driven approach uncovers unrecognized metastatic patterns and provides an unbiased framework for identifying cancer-type-specific trends, thereby addressing a longstanding challenge in metastasis research.

To demonstrate the versatility and scalability of Metient, we applied it to a *KRAS*-mutant lung adenocarcinoma cell line with single-cell lineage tracing, a domain where migration history reconstruction was previously constrained by only simplified models of metastatic spread. In these data, Metient identified the mediastinum as a key hub for further metastatic dissemination after initial seeding from the lung primary, revealing the particular ordering of seeding that a previous simplified model was not able to infer. Unlike prior multi-objective approaches built on integer linear programming, Metient leverages gradient-based optimization that can efficiently utilize GPUs, enabling scalability to larger and more complex datasets. This capability opens new avenues for studying metastasis at single-cell resolution, offering a powerful tool for future research.

Currently, Metient uses genetic distance and organotropism as its metastasis priors, as well as MACHINA’s parsimony criteria, however, the Metient framework is designed to be easily extensible. New parsimony criteria can be added as long as they can be computed as matrix functions of the tree adjacency matrix and vertex labeling matrices. Adding a new prior simply requires writing a scoring function because Metient incorporates auto-differentiation to compute its gradient updates. For instance, the framework could be easily extended to incorporate mutational signatures as a prior, since metastases exhibit shifts in mutational signature composition ^53,54^. Metient has some limitations. It scales well in compute time for larger clone trees or more samples but, because the loss landscape complexity increases substantially, in approximately 1% of cases in bulk-sequencing data and 5% of single-cell lineage tracing data, Metient became stuck in local minima. This problem was resolved when we ran Metient multiple times and with more samples, and we recommend this practice with larger reconstruction problems. One criteria to assess convergence is when the Pareto front remains unchanged. Other migration history algorithms are also highly sensitive to the complexity of the loss landscape, and convergence issues that they face are not necessarily resolved by rerunning the algorithm. Also, Metient is not designed to consider subclonal copy number alternations (CNAs) when correcting its estimated variant allele frequencies for CNAs. Using the descendant cell fraction (DCF) ^55^ or phylogenetic cancer cell fraction (phyloCCF) ^56^ as inputs to Metient could solve this. Alternatively, one could input which clones are in which samples directly into Metient instead of the allele frequencies that define subclones, as we did to run Metient on the lineage tracing data. Finally, we note that choice of clustering and tree inference algorithm used when inputting data into Metient can impact both the clonality and phyleticity classifications. In an attempt to most accurately compare our migration histories to previously reported results, where possible, we used the same clustering and trees inferred for the original datasets.

In conclusion, we show that Metient offers a fast and adaptable, fully automated framework that leverages molecular tumor sequencing data to probe enduring questions in metastasis research.

## Methods

### Estimating observed clone proportions

The first step of Metient is to estimate the binary presence or absence of clone tree (**T**) nodes in each site. The clone tree **T** can either be provided as input, or inferred from the DNA sequencing data using, e.g., Orchard ^57^, PairTree ^58^, SPRUCE ^59^, CITUP ^60^, or EXACT ^61^. Building on a previous approach as described by Wintersinger et al. ^58^, Metient estimates the proportion of clones in each site using the input clone tree **T** and read count data from bulk DNA sequencing. For a genomic locus *j* in anatomical site *k*, the probability of observing read count data *x*_*kj*_ is defined using the following:

- *A*_*kj*_ is the number of reads that map to genomic locus *j* in anatomical site *k* with the variant allele
- *R*_*kj*_ is the number of reads that map to genomic locus *j* in anatomical site *k* with the reference allele
- *ω*_*kj*_ is a conversion factor from mutation cellular frequency to variant allele frequency (VAF) for genomic locus *j* in anatomical site *k*

Using a binomial model, we then estimate the proportion of anatomical site *k* containing clone *c* using *p*(*x*_*kj*_| **F**_*kj*_)= Binom(*A*_*kj*_|*A*_*kj*_ + *R*_*kj*_, *ω*_*kj*_ **F**_*kj*_). Where **F** = **UB** is the mutation cellular frequency matrix, **B** ∈ {0, 1}^*C*×*M*^ is 1:1 with a clone tree, where *C* is the number of clones and *M* is the number of mutations or mutation clusters, and **B**_*cm*_ =1 if clone *c* contains mutation *m* (Figure S1b). **U** ∈ [0, 1]^*K*×*C*^, where *K* is the number of anatomical sites, and **U**_*kc*_ is the fraction of anatomical site *k* made up by clone *c* (Figure S1b). An L1 regularization is used to promote sparsity, since we expect most values in **U** to be zero. For details on how to set *ω*_*kj*_, see “Variant read probability calculation (*ω*)” in Supplementary Information. An alternative way to find a point estimate of **U** is using a previously described projection algorithm for this problem ^57,58,61,62^. A point estimate **U** can be found by optimizing the following quadratic approximation to the binomial likelihood of **U** given **B** and **F**:

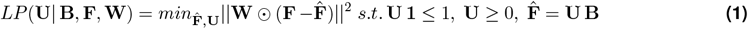

where ||·|| is the Frobenius norm, **1** is a vector of 1s, **F** are the observed mutation frequencies, **W** is a *K* × *M* matrix of inverse-variances for each mutation in each sample derived from **F**, and ⊙ is the Hadamard, i.e., element-wise product. The definition for **W** is as described in previous work ^58,61^.

We use **U** (estimated in either of the previously described ways) to determine if a clone *c* is present in an anatomical site *k*. If *c* is present, we attach a witness node with label *k* (leaf nodes connected by dashed lines in Figure S1b, c) to clone *c* in clone tree **T**. We deem *c* to be present in *k* if **U**_*kc*_ > 5% for a given anatomical site *k* and clone *c*. If a clone *c* does not make up 5% of any of the *K* anatomical sites, and *c* is a leaf node of the clone tree **T**, we remove this node since it is not well estimated by the data.

Here the term “anatomical site” is used to describe a distinct tumor mass. If multiple samples are taken from the same tumor mass, we combine them as described in “Bulk DNA sequencing pre-processing: Non-small Cell Lung Cancer Dataset”.

Note that read count data are only used to determine which clones are present in which sites, if a matrix indicating the presence or absence of each clone in each anatomical site is available, it can be used as an input to replace the read count data. These clone-to-site assignment matrices can be derived, e.g., from single-cell data.

### Labeling the clone tree

The next step in inferring a migration history is to jointly infer a labeling of the clone tree and resolve polytomies, i.e., nodes with more than two children. Polytomy resolution is discussed in the section “Resolving polytomies”.

Because we are interested in identifying multiple hypotheses of metastatic spread, Metient seeks to find multiple possible labelings of a clone tree **T**. Each possible labeling is represented by a matrix **V** ∈ {0, 1}^*K*×*C*^, where *K* is the number of anatomical sites and *C* is the number of clones, and **V**_*kc*_ =1 if clone *c* is first detected in anatomical site *k*. Each column of **V** is a one-hot vector. We solve for an individual **V** by optimizing the evidence lower bound, or ELBO, as defined by:

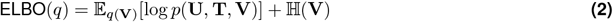

Where 𝔼_*q*(**V**)_[log*p*(**U, T, V**)] evaluates a labeling based on parsimony, genetic distance, and organotropism, and the second term is the entropy term. **U** has been optimized as described in the previous section “Estimating observed clone proportions”, or taken as input from the user. See Supplementary Information for a full derivation of this objective. Because **V** is a matrix of discrete categorical variables, we do not optimize **V** directly, but rather the underlying probabilites of each category that we optimize using a Gumbel-softmax estimator (see “Gumbel-softmax optimization”).

### Gumbel-softmax optimization

In the previous section, we described how to score the matrix representation of the labeled clone tree, **V**. Here, we describe how to optimize **V** via the straight-through estimator of the Gumbel-Softmax distribution ^63,64^. Starting with a matrix *ψ* ∈ {0, 1}^*K*×*C*^, of randomly initialized values, where *K* is the number of anatomical sites and *C* is the number of clones, and each column represents the unnormalized log probabilities of clone *c* being labeled in site *k*:

1. At every iteration, for each clone *c*, we sample *g*_1*c*_…*g*_*kc*_, *k* i.i.d. samples from Gumbel(0,1) and compute *y*_*ic*_ = *ψ* _*ic*_ + *g*_*ic*_.
2. We then sample from the categorical distribution represented by the column vector *ψ*_:*c*_ by setting *i*^*^ = argmax_*i*_ *y*_*ic*_ and represent that sample with a one-hot encoding in **V**, i.e., **V**_*ic*_ =1 if *i* = *i*^*^, 0 otherwise.
3. Then we evaluate the ELBO(*ν*) where

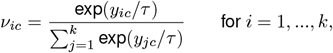

using a stochastic approximation based on **V**, and take the gradient of this ELBO in the backward pass, thus implementing the straight-through estimator.
4. During training, start with a high *τ* to permit exploration, then gradually anneal *τ* to a small but non-zero value so that the Gumbel-Softmax distribution, *ν* resembles a one-hot vector.

At the end of training, as *τ* approaches 0, then the gradient becomes unbiased and *ν* approaches **V**. In order to capture multiple modes of the posterior distribution, each representing different hypotheses about the migration history, we optimize multiple **V**s in parallel. To do this, we set up steps 1-3 such that *x ψ*s are solved for in parallel ^65^ (with a different random initialization for each parallel process), where *x* is equal to the sample size and is calculated according to the size of the inputs (α *K*^*C*^). See Supplementary Information for further explanation.

### Resolving polytomies

An overview of the algorithm to resolve polytomies is given in Supplementary Figure S11a and b.

1. If a node *i* in **T** has more than 2 children, we create *r* new “resolver” nodes where *r* is the number of children of node *i/*2. These new resolver nodes’ vertex labels are jointly inferred along with the refinement of **T**. The genetic distance between the parent node *i* and its new resolver node is set to 0 since there are no observed mutations between the two nodes.
2. We allow the children of *i* to stay as a child of *i*, or become a child of one of the resolver nodes of *i*.
3. Any resolver nodes that are unused (i.e. have no children) or which do not improve the migration history (i.e. the parsimony metrics without the resolver node are the same or worse) are removed.

### Fixing optimal subtrees

To improve convergence, we perform two rounds of optimization when solving for a labeled clone tree and resolving polytomies:

1. Solve for labeled trees and resolve polytomies jointly (as described in previous sections).
2. For each pair of labeled tree and polytomy resovled tree, find optimal subtrees. I.e., find the largest subtrees, as defined by the most number of nodes, where all labels for all nodes are equal. This means that there is no other possible optimal labeling for this subtree (there are 0 migrations, 0 comigrations, 0 seeding sites), and we can keep it fixed. Fix these nodes’ labelings and adjacency matrix connections (if using polytomy resolution).
3. Repeat step 1 for any nodes that have not been fixed in step 2.

### Guaranteeing minimum migration solutions on the Pareto front

To guarantee a minimum migration solution on the Pareto front, when polytomy resolution is not being used, we run the Fitch-Hartigan algorithm ^20,21^. This aids in output stability when input sizes are very large, however Metient recovers these solutions using its sampling approach (along with multiple minimum migration solutions) in almost all cases.

### Metient-calibrate

In Metient-calibrate, we aim to fit a patient cohort-specific parsimony model using the metastasis priors. To score a migration history using genetic distance, we use the following equation: Σ_*ij*_ −*log*(**D**_**ij**_)**K**_*ij*_, where **D** contains the normalized number of mutations between clones, and **K** =1 if clone *i* is the parent of clone *j* and clone *i* and clone *j* have different anatomical site labels.

To score a migration history using organotropism, we use the following equation: 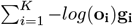, where vector **o** contains the frequency at which the primary seeds other anatomical sites, and vector **g** contains the number of migrations from the primary site to all other anatomical sites for a particular migration history.

To optimize the parsimony metric weights, Metient identifies a Pareto front of labeled trees for each patient and scores these trees based on (1) the weighted parsimony metrics and (2) the metastasis priors: genetic distance and, if appropriate anatomical labels are available, organotropism. These form the parsimony distribution and metastasis prior distribution, respectively. We initialize with equal weights and use gradient descent to minimize the cross entropy loss between the parsimony distribution and metastasis prior distribution for all patients in the cohort. Once the optimization converges, Metient rescores the trees on the Pareto front using the fitted weights, to identify the maximum calibrated parsimony solution, and genetic distance and organotropism are used to break ties between equally parsimonious migration histories. See Supplementary Information for a more detailed derivation.

### Metient-evaluate

In Metient-evaluate, weights for each maximum parsimony metric (migrations, comigrations, seeding sites) and optionally, genetic distance and organotropism, are taken as input. These weights are used to rank the solutions on the Pareto front. If no weights are inputted, we provide a pan-cancer parsimony model calibrated to the four cohorts (melanoma, HGSOC, HR-NB, NSCLC) discussed in this work.

### Defining the organotropism matrix

Data from the MSK-MET study ^31^ for 25,775 patients with annotations of distant metastases locations was downloaded from the publicly available cbioportal ^66^. Each patient had annotations of one of 27 primary cancer types and the presence or absence of a metastasis in one of 21 distant anatomical sites. The original authors extracted this data from electronic health records and mapped it to a reference set of anatomical sites. We sum over all patients to build a 27 x 21, cancer type by metastatic site occurrence matrix. We then normalize the rows to turn these into frequencies. We interpret the negative log frequencies as a “relative time to metastasis”, and only score migrations from the primary site to other sites, because there is no data to indicate frequencies of seeding from metastatic sites to other metastatic sites, or back to the primary. We make this data available for users, with the option for users to instead input their own organotropism vector for each patient.

### Evaluations on simulated data

We use the simulated data for 80 patients provided by MACHINA ^17^ to benchmark our method’s performance. To prepare inputs to Metient, we use the same clustering algorithm and clone tree inference algorithm used in MACHINA (MACHINA ^17^ and SPRUCE ^59^, respectively) in order to accurately compare only our migration history inference algorithm (including polytomy resolution) against MACHINA’s. All performance scores are reported using MACHINA’s PMH-TI mode and Metient-calibrate with a sample size of 1024, both with default configurations. We do not use polytomy resolution for Metient-calibrate in these results, since it does not improve performance on simulated data. (Supplementary Tables 4, 3). However, this performance is not necessarily indicative of polytomy resolution working poorly, because it actually finds more parsimonious solutions than the ground truth solution in 75% of simulated data (Supplementary Figure S10).

#### Evaluation metrics

We use the same migration graph and seeding clones F1-scores as MACHINA. Given a reconstructed migration graph **G**, its recall and precision with respect to the ground truth migration graph **G**^∗^ are calculated as follows:

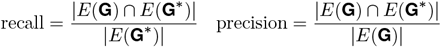

where *E*(**G**) are the edges of **G**, and multiple edges between the same two sites are included in *E*(**G**). When there are multiple edges from site *i* to site *j*, |*E*(**G**) ∩ *E*(**G**^∗^)| = min(*a, b*), where *a* and *b* are the number of edges from site *i* to site *j* in **G** and **G**^∗^, respectively.

Recall and precision of the seeding clones in the inferred migration history (which includes inference of both the clone tree labeling and observed clone proportions) is calculated as follows:

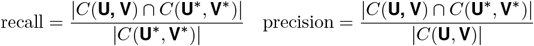

where *C*(**U, V**) is the set of mutations, i.e., the subclone, associated with the clone nodes that have an outgoing migration edge. For example, *C*(**U, V**) = A, B, C in solution A of Figure S1c. The definition for seeding clones used in these evaluations is distinct from how we define seeding clones in the rest of the paper (“Defining colonizing clones, clonality, and phyleticity” in Methods). Specifically, if there is an edge between two nodes (*u, υ*), where the labeling of *u* and *υ* are not equal, we define the seeding clone as *υ*. However in order to consistently compare to MACHINA in these evaluations, we use their definition and define the seeding clone as *u*. We note that identifying the mutations of *υ* is generally a harder problem.

#### Timing benchmarks

All timing benchmarks (Figure S3e) were run on 8 Intel(R) Xeon(R) CPU E5-2697 v4 @ 2.30GHz CPU cores with 8 gigabytes of RAM per core. Runtime of each method is the time needed to run inference and save dot files of the inferred migration histories (and for Metient, an additional serialized file with the results of the top k migration histories). We compare MACHINA’s PMH-TI mode to Metient-calibrate with a sample size of 1024, both with default configurations. These are the same modes used to report comparisons in F1-scores. Each value in Figure S3e is the time needed to run one patient’s tree. Because Metient-calibrate has an additional inference step where parsimony metric weights are fit to a cohort, we take the time needed for this additional step and divide it by the number of patient trees in the cohort, and add this time to each patient’s migration history runtime.

### Defining colonizing clones, clonality, and phyleticity

A colonizing clone is defined as a node in a migration history whose parent is a different color than itself. There are two exceptions to this rule: when node *a* has a parent with a different color than itself, but the node is a witness node (Figure S1c) or a polytomy resolver node (e.g. A_POL in Supplementary Figure S11a). In these cases, these nodes do not represent any new mutations, but rather contain the same mutations as its parent. For these two cases, the colonizing clone is defined to be *a*’s parent node.

In order to rectify different meanings of the terms “monoclonal” and “polyclonal” used in previous work, we define two terms:

- genetic clonality: if all sites are seeded by the same colonizing clone, this patient is genetically monoclonal, otherwise, genetically polyclonal.
- site clonality: if each site is seeded by one colonizing clone, but not necessarily the same colonizing clone, this patient is site monoclonal, otherwise, site polyclonal.

Genetic clonality and site clonality are depicted schematically in Figure 2b.

To define phyleticity, we first extract all colonizing clones from a migration history. We then identify the colonizing clone closest to the root, *s*, i.e., the colonizing clone with the shortest path to the root. If all other colonizing clones are descendants of the tree rooted at *s*, the migration history is monophyletic, otherwise, it is polyphyletic. Under this definition, if a tree is monophyletic, then there are no independent evolutionary trajectories that give rise to colonizing clones. This is depicted schematically in Figure 2c.

In order to accurately compare our phyleticity measurements to TRACERx, we use their definition in Figure 4c and the TRACERx comparison analysis. To apply their definition to our migration histories, we extract colonizing clones as described above, and then determine if there is a Hamiltonian path in the clone tree that connects the colonizing clones. I.e., we determine if there is a path in the clone tree that visits each colonizing clone exactly once. If such a Hamiltonian path exists, we call this migration history monophyletic under the TRACERx definition, and polyphyletic otherwise.

### Bootstrap sampling for fitting parsimony metric weights

Running Metient-calibrate on the 167 patients from the melanoma, HGSOC, HR-NB and NSCLC datasets infers a Pareto front of migration histories for each patient. For each dataset, we subset patients that have a Pareto front with size greater than one, and take 100 bootstrap samples of patients from this subset. Patients with a single solution on the Pareto front do not have an impact on the cross-entropy loss used to fit the parsimony metric weights. For each bootstrap sample of patients, their Pareto front migration histories are used to fit the parsimony metric weights (“Calibrate alignment” in Supplementary Information). For each of the parsimony metric weights fit to a bootstrap sample, we evaluated how these weights would order the Pareto front, and evaluated how consistently the same top solution was chosen. We average the percent of times the same solution is ranked as the top solution across the four datasets.

## Data availability

The results for the HR-NB cohort were based on data under the study phs03111.v1.p1 and were accessed from the NCI’s Cancer Research Data Commons (https://datacommons.cancer.gov). The anatomical site labels in Figure 6 used data generated by the TRAcking Non-small Cell Lung Cancer Evolution Through Therapy (Rx) (TRACERx) Consortium and provided by the UCL Cancer Institute and The Francis Crick Institute. The TRACERx study is sponsored by University College London, funded by Cancer Research UK and coordinated through the Cancer Research UK and UCL Cancer Trials Centre. This data is available from the European Genome-phenome Archive under accession EGAD00001009825^67^. All other data used for the NSCLC cohort were made publicly available and accessed at https://zenodo.org/records/7649257. The organotropism matrix derived from data in the MSK-MET study ^31^ is available at https://github.com/morrislab/metient/blob/main/metient/data/msk_met/msk_met_freq_by_cancer_type.csv. The following publicly available datasets were accessed via the supplementary information in the following studies: melanoma ^3^, breast ^68^, HGSOC ^4^, MSK-MET ^31^. The lung adenocarcinoma single-cell lineage tracing data were made publicly available and accessed at https://zenodo.org/records/4243162.

## Code availability

Metient is available as a software package installable with pip at https://github.com/morrislab/metient/. Tutorials for usage can be found at https://github.com/morrislab/metient/tree/main/tutorial. Code to reproduce figures from this manuscript can be found at https://github.com/morrislab/metient/tree/main/metient/jupyter_notebooks.

## Acknowledgments

We thank Julia Simundza for her valuable feedback on this manuscript, and Deeksha Madala for coming up with the method name Metient. K.G. is supported by NIH grants R37CA266185, U2CCA233284 and U54CA274492. This material is based upon work supported by the National Science Foundation Graduate Research Fellowship under Grant No. 227260-01 (D.K.) and NIH/NCI Cancer Center Support Grant P30 CA008748 (Vickers).

## Supplementary Information

**Figure S1.**
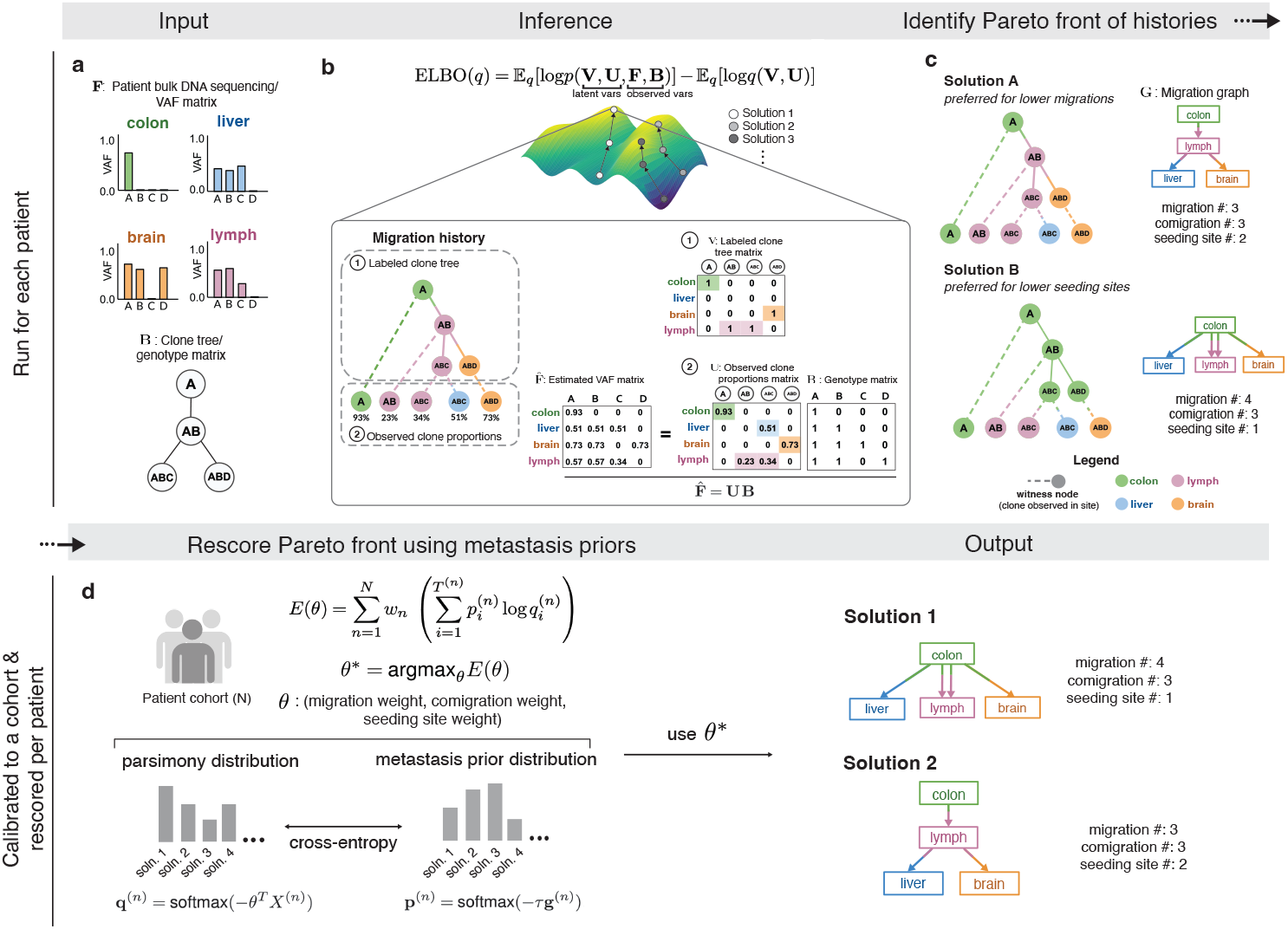
The Metient algorithm. **(a) Input**: (top) bulk DNA sequencing sampled from multiple tumors in a single patient, and (bottom) a clone tree which represents the evolutionary relationship of mutations. AB refers to a clone with mutations or mutation clusters A and B. **(b) Inference**: Using the inputs as observed variables, we infer the latent variables (1) **V** (representing the labeled clone tree) and (2) **U** (representing the proportion of each clone in each anatomical site). 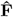 is the estimated VAF matrix produced by **UB**, where **B**_*ij*_ = 1 if clone *i* contains mutation *j*. Each migration history solution can be represented by a migration history, which is a clone tree with (1) an anatomical site labeling of its internal nodes, and (2) leaf nodes representing the observed clone proportions in anatomical sites. **(c) Identify Pareto front of histories**: We infer a Pareto front of migration histories as defined by the three parsimony metrics (migration, comigration and seeding site number). A migration graph **G** summarizes the migration edges of the migration history. **(d) Metient-calibrate**: Weights on the parsimony metrics (*θ*) are fit by minimizing the cross entropy loss between each patient’s migration histories’ probability distribution as scored by the metastasis priors (target distribution) and the probability distribution as scored by the parsimony metrics (source distribution). These weights are fit across a cohort of patients, and then used to rescore the Pareto front of migration histories produced for each patient in that cohort.

**Table 1.** Summary of previous methods which perform some aspect of migration history inference. Y = yes, N = no. Labels clone tree refers to whether the method infers the labels of the internal vertices of a clone tree (e.g. labeling clone AB as originating in lymph in Figure S1c, solution A). Estimates clone proportions in sites refers to whether the method infers the leaf nodes (witness nodes) (e.g. identifying that clone ABC is present in both lymph and liver in Figure S1c, solution A). Multiple solutions indicates whether a method outputs multiple possible migration histories.

| Method | Previous Methods for Migration History Inference |  |  |  |  |  |  |
| --- | --- | --- | --- | --- | --- | --- | --- |
|  | Labels clone tree | Estimates clone proportions in sites | Models Complex Seeding | Multiple solutions | Organo-tropism | Genetic Distance | Polytomy Resolution |
| ClonEvol<sup>15</sup> | Y | Y | N | Y | N | N | N |
| Treeomics<sup>16</sup> | N | Y | N | N | N | N | N |
| MACHINA<sup>17</sup> | Y | Y | Y | N | N | N | Y |
| PathFinder<sup>18</sup> | Y | N | N | Y | N | Y | Y |
| Metient | Y | Y | Y | Y | Y | Y | Y |

**Table 2.**
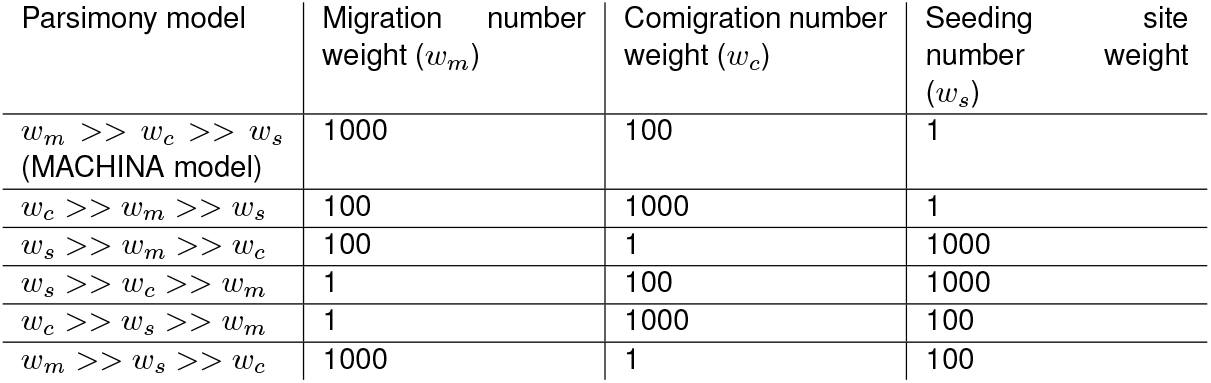
The multiple parsimony models that Metient uses to build a Pareto front of solutions for a patient’s data. Each parsimony model has a different relative weighting on each parsimony metric.

**Figure S2.**
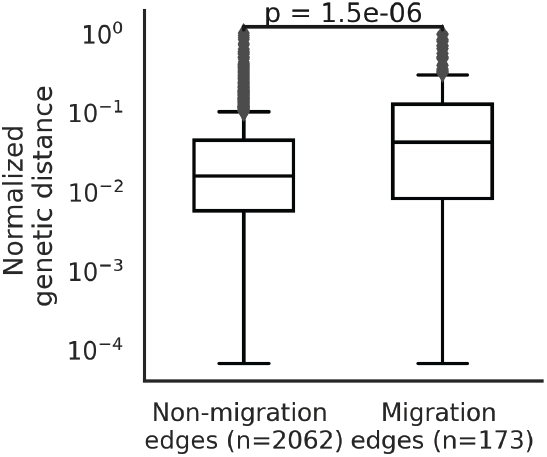
In 92 patients with a single migration history on the Pareto front, the distribution of normalized genetic distance on non-migration edges vs. migration edges. Normalized genetic distance for a branch of a clone tree is calculated as the number of mutations on a branch divided by the maximum number of mutations on any branch of the clone tree. Statistical significance assessed by a Welch’s t-test.

**Figure S3.**
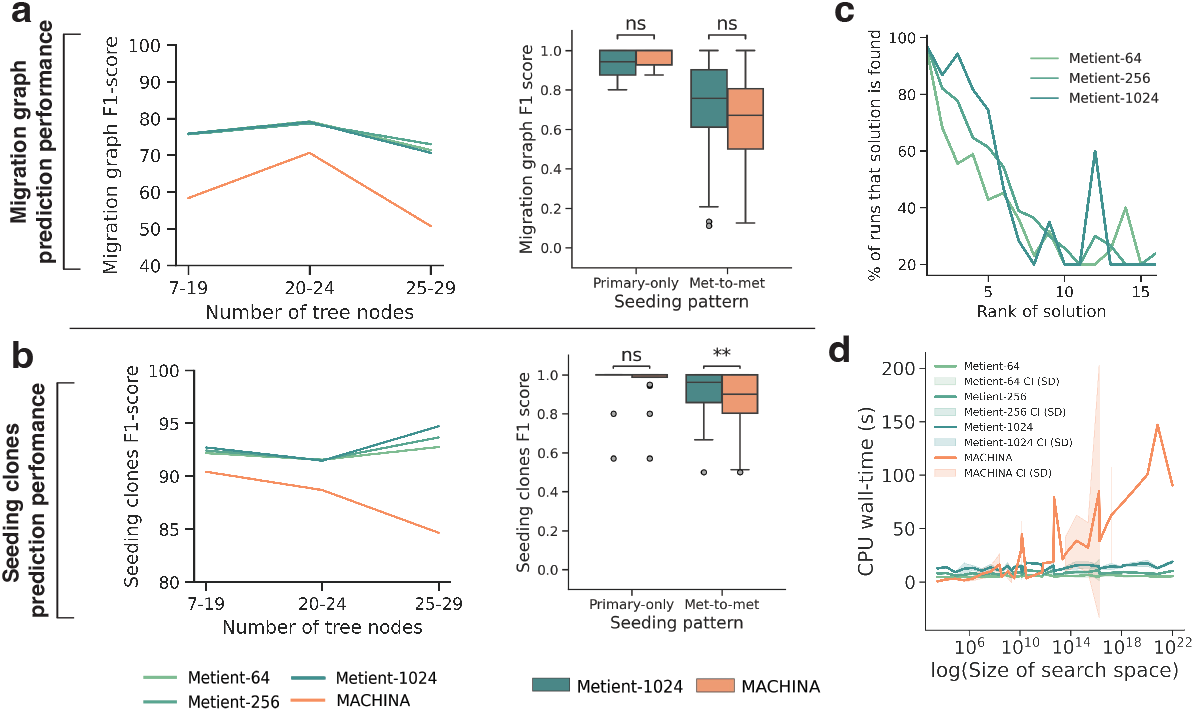
Metient achieves state-of-the-art performance on simulated data. All results shown for Metient are in calibrate mode using genetic distance as the metastasis prior. Metient-1024 refers to a model configuration where 1024 solutions are sampled. For a given simulated input, for MACHINA (which outputs one solution) the top solution is used, and for Metient we evaluate all top (lowest loss) solutions. **(a)** The averaged F1-score for predicted the ground-truth migration graph (left), and the distribution of F1-scores (right) for patients with primary-only seeding (n=16) vs. patients with metastasis-to-metastasis (met-to-met) seeding (n=64). Statistical significance assessed by a Wilcoxon signed rank test; ns: not significant. **(b)** The averaged F1-score for predicted the ground-truth seeding clones (left), and the distribution of F1-scores (right) for patients with primary-only seeding (n=16) vs. patients with metastasis-to-metastasis (met-to-met) seeding (n=64). Statistical significance assessed by a Wilcoxon signed rank test; ns: not significant, **: p=0.0021. **(c)** After running Metient five times, the percentage of runs that a certain solution is found as a function of its averaged rank across runs. **(d)** CPU wall-time needed to run Metient vs. MACHINA as a function of the search space size. CI: confidence interval, SD: standard deviation.

**Table 3.** Average F1-scores of migration graph for each broad seeding pattern (primary-only seeding or metastasis-to-metastasis seeding) on simulated data. All Metient models were run with a sample size of 1024. When multiple solutions are found for a given input, all lowest loss solutions were taken. Evaluate (MP): Metient in evaluate mode with maximum parsimony only. Evaluate (GD): Metient in evaluate mode with genetic distance only. Calibrate: Metient in calibrate mode, using genetic distance as the metastasis prior. polyres: polytomy resolution is used. mS: monoclonal single-source seeding. pS: polyclonal single-source seeding. pM: polyclonal multi-source seeding. pR: polyclonal reseeding.

**Average migration graph F1-scores**
| Method | Primary-only | Met-to-met | Macro-F1 | Micro-F1 |
| --- | --- | --- | --- | --- |
| Evaluate (MP) | 0.930 | 0.688 | 0.809 | 0.736 |
| Evaluate (MP) + polyres | **0.983** | 0.648 | 0.816 | 0.715 |
| Evaluate (GD) | 0.857 | 0.691 | 0.774 | 0.724 |
| Evaluate (GD) + polyres | 0.829 | 0.649 | 0.739 | 0.685 |
| Calibrate | 0.930 | **0.716** | **0.823** | **0.759** |
| Calibrate + polyres | **0.983** | 0.662 | **0.823** | 0.726 |
| MACHINA | 0.968 | 0.643 | 0.806 | 0.708 |

**Table 4.** Average F1-scores of migrating clones for each broad seeding pattern (primary-only seeding or metastasis-to-metastasis seeding) on simulated data. All Metient models were run with a sample size of 1024. When multiple solutions are found for a given input, all lowest loss solutions were taken. Evaluate (MP): Metient in evaluate mode with maximum parsimony only. Evaluate (GD): Metient in evaluate mode with genetic distance only. Calibrate: Metient in calibrate mode, using genetic distance as the metastasis prior. polyres: polytomy resolution is used.

| Method | Primary-only | Met-to-met | Macro-F1 | Micro-F1 |
| --- | --- | --- | --- | --- |
| Evaluate (MP) | 0.795 | 0.781 | 0.788 | 0.784 |
| Evaluate (MP) + polyres | 0.873 | 0.791 | 0.832 | 0.808 |
| Evaluate (GD) | 0.954 | 0.876 | 0.915 | 0.892 |
| Evaluate (GD) + polyres | **0.979** | **0.928** | **0.954** | **0.939** |
| Calibrate | 0.961 | 0.916 | 0.938 | 0.925 |
| Calibrate + polyres | 0.961 | 0.890 | 0.926 | 0.905 |
| MACHINA | 0.954 | 0.876 | 0.915 | 0.892 |

**Figure S4.**
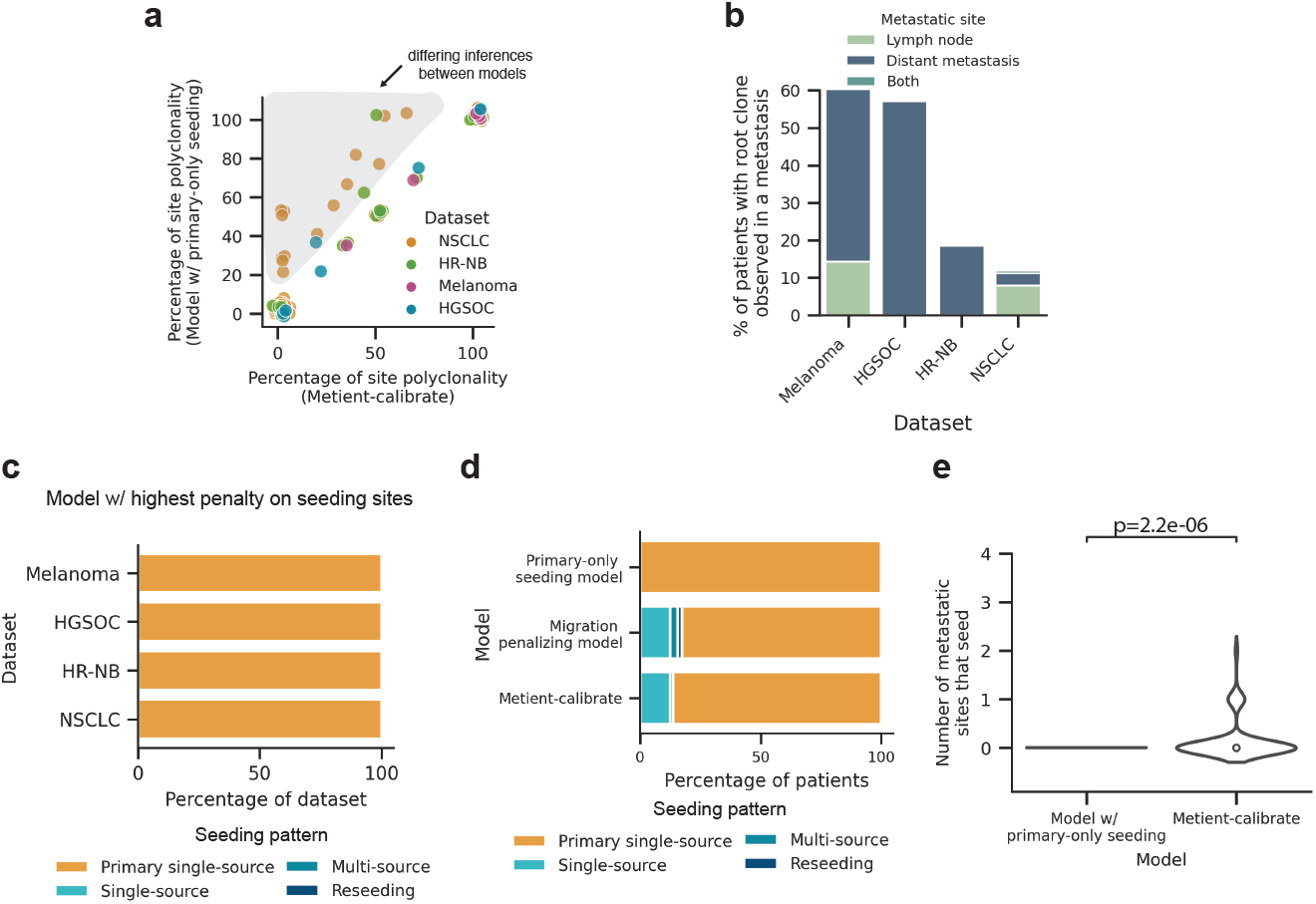
**(a)** A comparison of the percent of site polyclonal migrations for each patient’s migration history when using the best migration history chosen by Metient (x-axis) vs. a model that assumes primary-only seeding (y-axis). **(b)** Percent of patients in each dataset with the root cancerous clone observed in a metastatic site. **(c)** The distribution of seeding patterns in each dataset when taking the migration history on the approximate Pareto front with the lowest number of seeding sites, run with Metient-calibrate. **(d)** The distribution of seeding patterns across all patients if we choose the migration history on the Pareto front with the lowest number of seeding sites (primary-only seeding model), lowest number of migrations (migration penalizing model), or the top Metient-calibrate solution. **(e)** A comparison of the number of metastatic sites that seed other sites between migration histories chosen by a model which chooses the migration history that assumes primary-only seeding (n=167) vs. Metient (n=167). Statistical significance assessed by a paired t-test.

**Figure S5.**
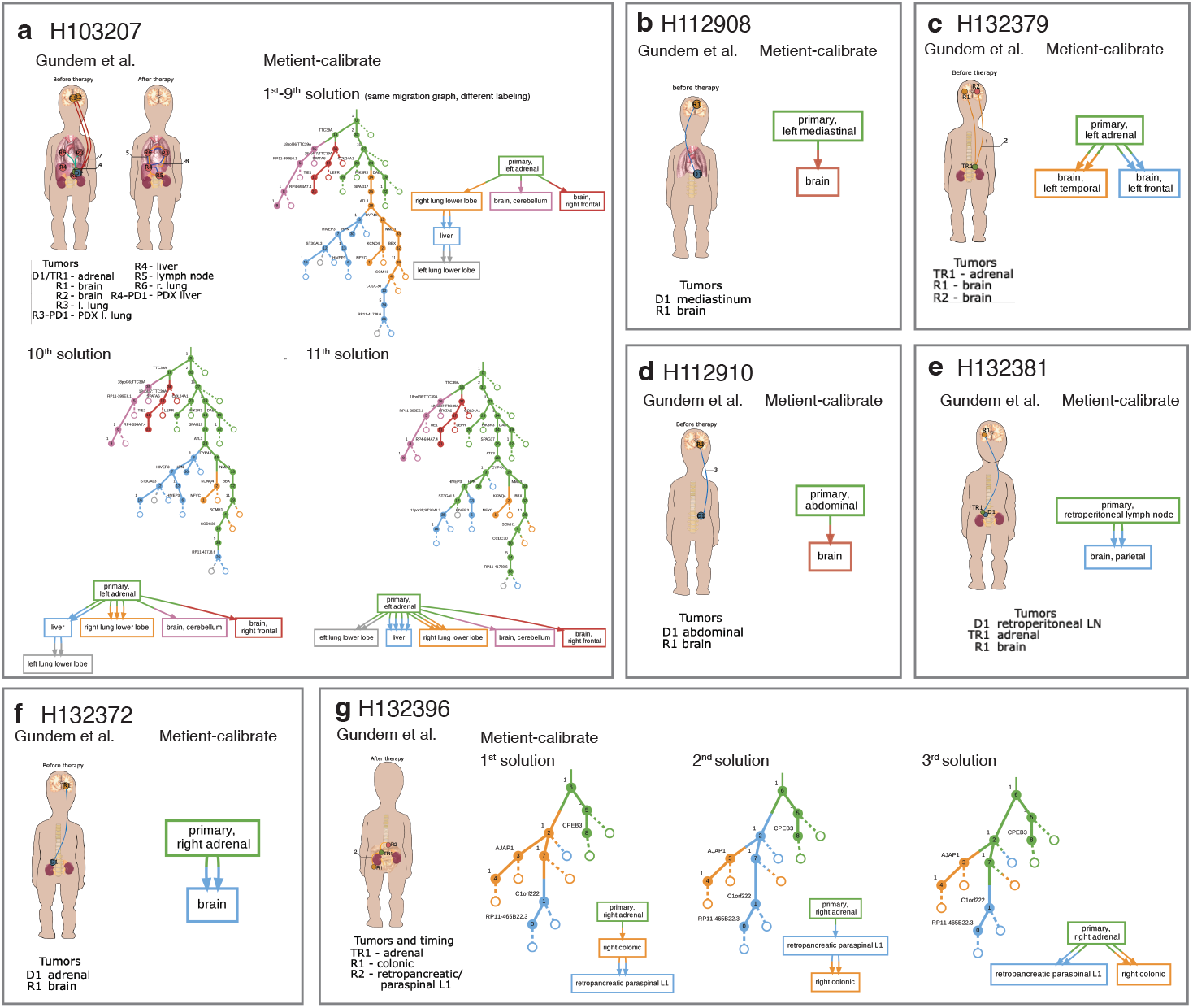
Comparison of Gundem et al. ^9^ reported body maps (left of each square) and Metient-calibrate inferred histories. The Metient-calibrate solutions with unique migration graphs on the Pareto front are shown. For example, in cases where there are multiple Pareto optimal migration histories with the same migration graph, only the migration history with the lowest loss is shown.

**Figure S6.**
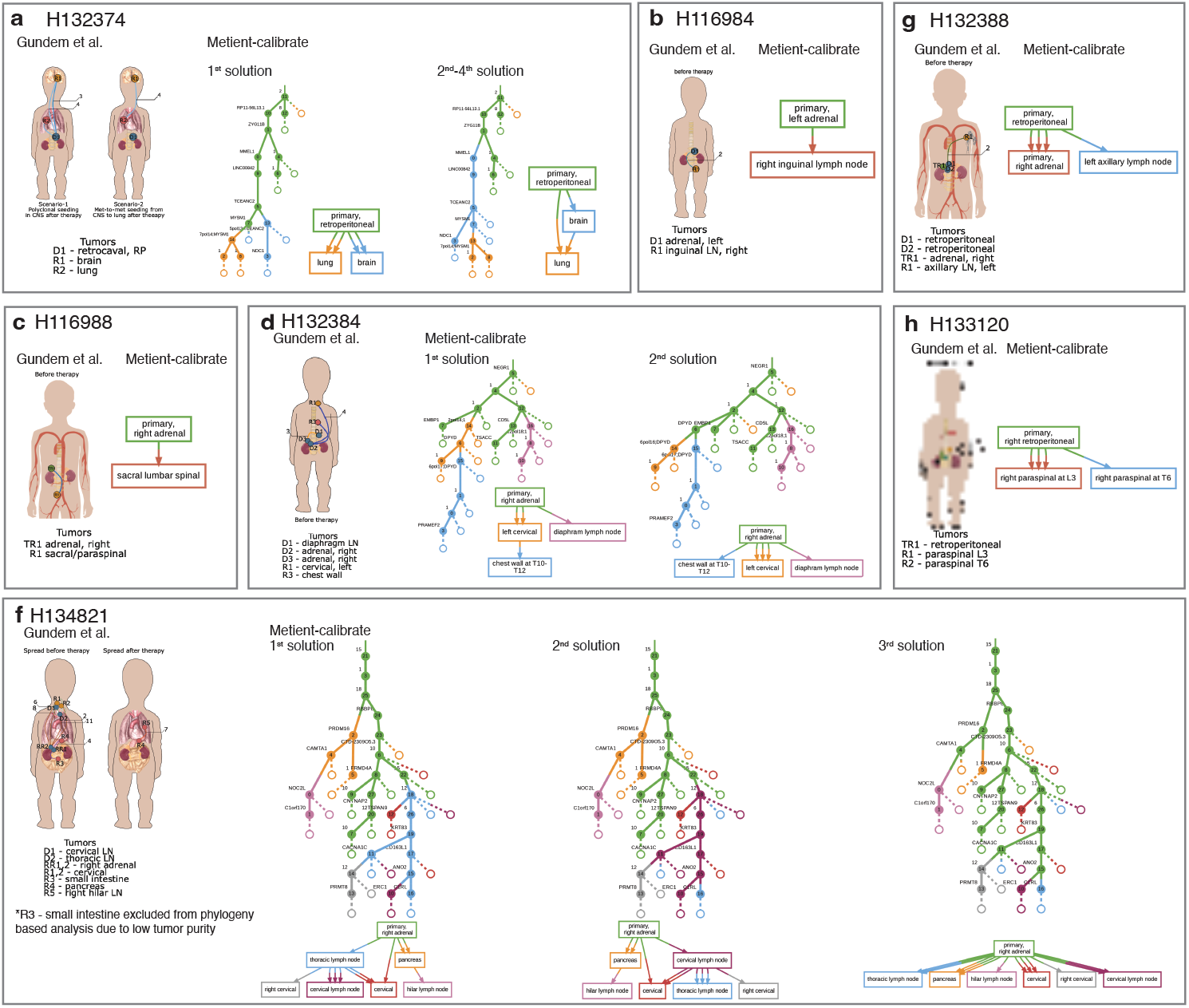
Comparison of Gundem et al. ^9^ reported body maps (left of each square) and Metient-calibrate inferred histories. The Metient-calibrate solutions with unique migration graphs on the Pareto front are shown. For example, in cases where there are multiple Pareto optimal migration histories with the same migration graph, only the migration history with the lowest loss is shown.

**Figure S7.**
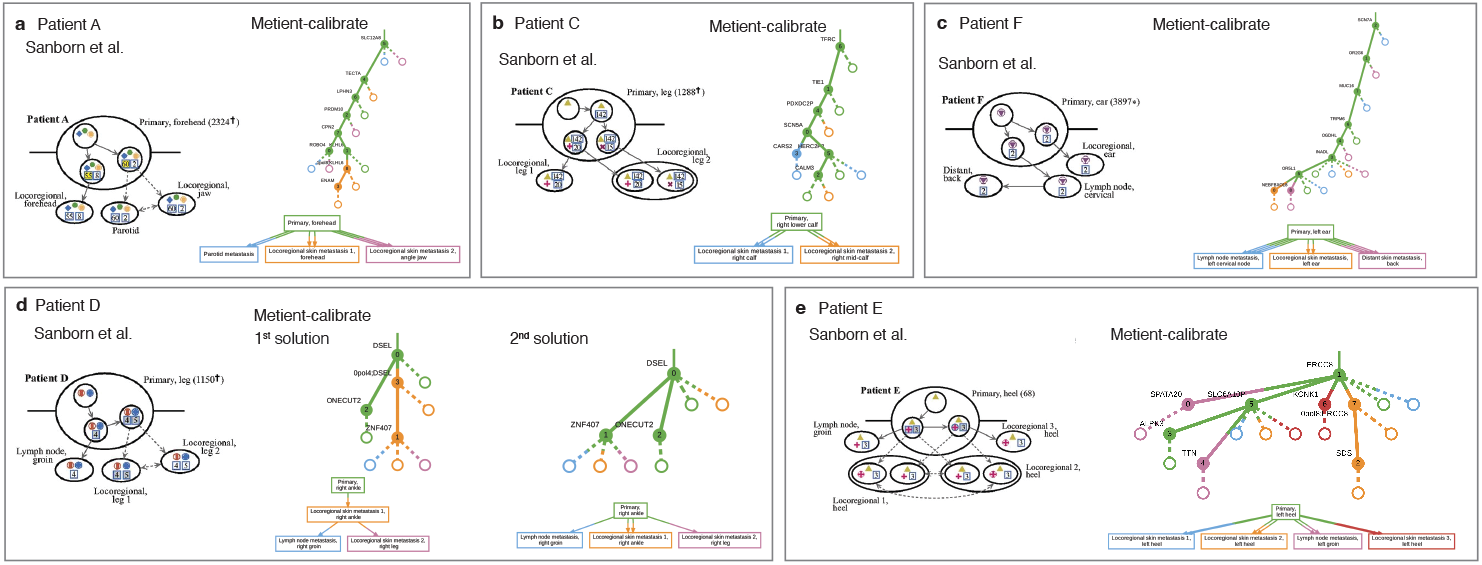
Comparison of Sanborn et al. ^3^ reported histories and Metient-calibrate inferred histories. In the Sanborn et al. ^3^ reported histories, solid lines denote probable dissemination patterns and dashed lines denote multiple possible paths. The Metient-calibrate solutions with unique migration graphs on the Pareto front are shown. For example, in cases where there are multiple Pareto optimal migration histories with the same migration graph, only the migration history with the lowest loss is shown.

**Figure S8.**
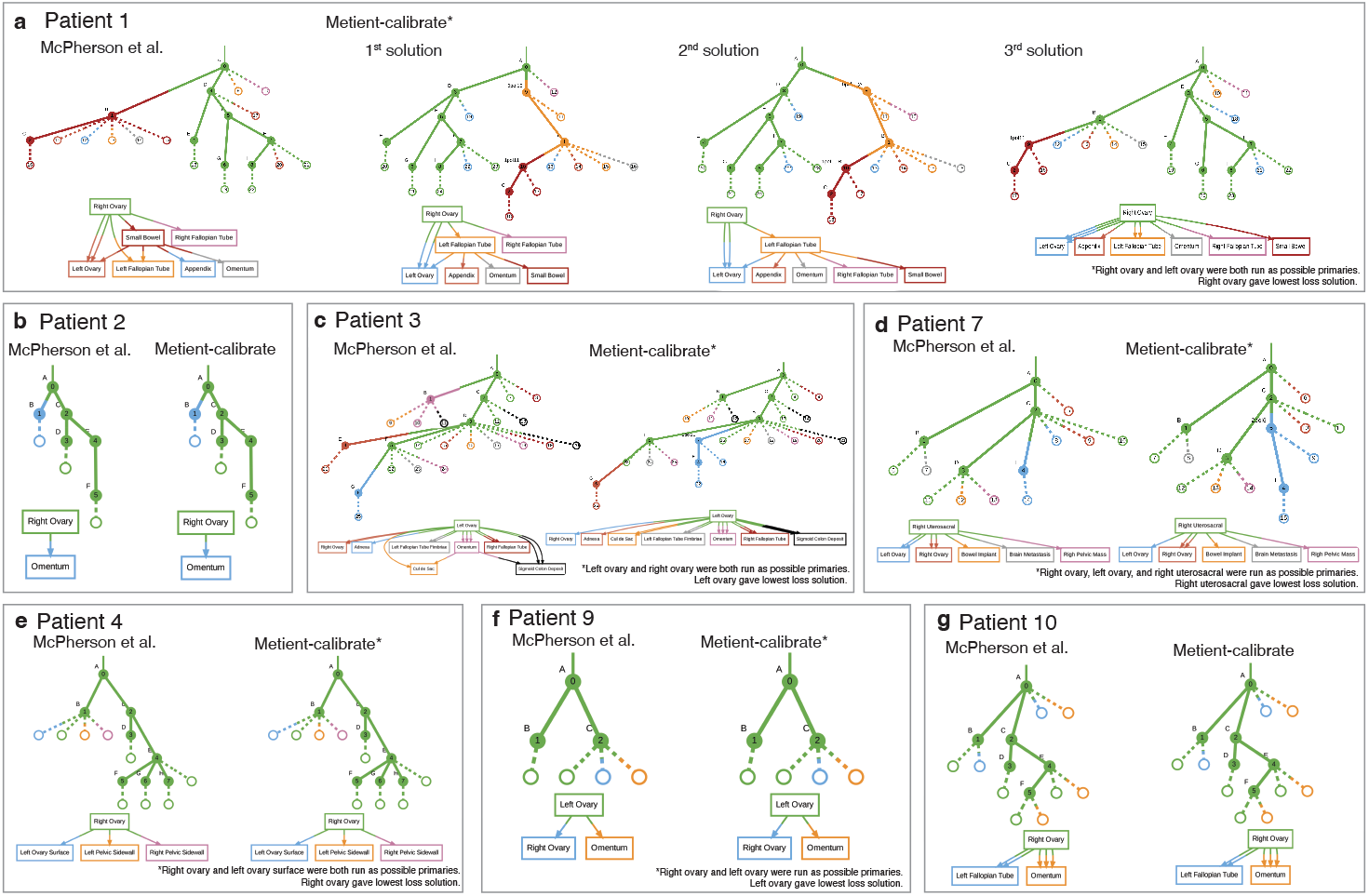
Comparison of McPherson et al. ^4^ reported histories and Metient-calibrate inferred histories. The Metient-calibrate solutions with unique migration graphs on the Pareto front are shown. For example, in cases where there are multiple Pareto optimal migration histories with the same migration graph, only the migration history with the lowest loss is shown. When multiple possible primaries were available, Metient-calibrate was run once with each possible primary, and the primary with the lowest loss solution is shown.

**Figure S9.**
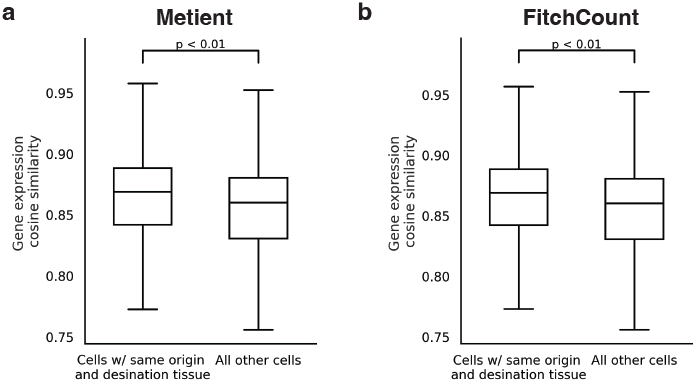
Cosine similarity between a cell’s gene expression profile and the averaged gene expression of cells from the same clone, either in Metient’s predicted site of origin or its destination (measured empirically) (1), or in a different tissue (2), using single-cell lineage tracing data ^22^. Statistical significance was assessed with a paired t-test (p = 1.3e-5).

**Figure S10.**
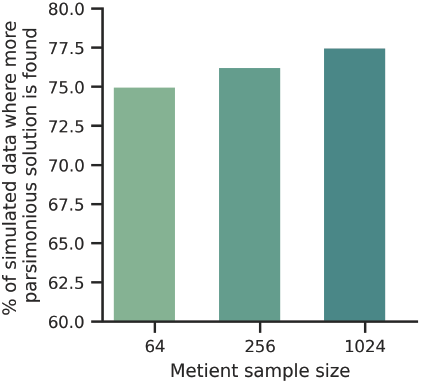
The percent of simulated data where a more parsimonious solution than ground truth is found when running Metient-1024 in calibrate mode with polytomy resolution. More parsimonious is defined as all of the parsimony metrics (migration, comigration and seeding site number) being less than or equal to the ground truth, but with at least one being strictly less than.

**Figure S11.**
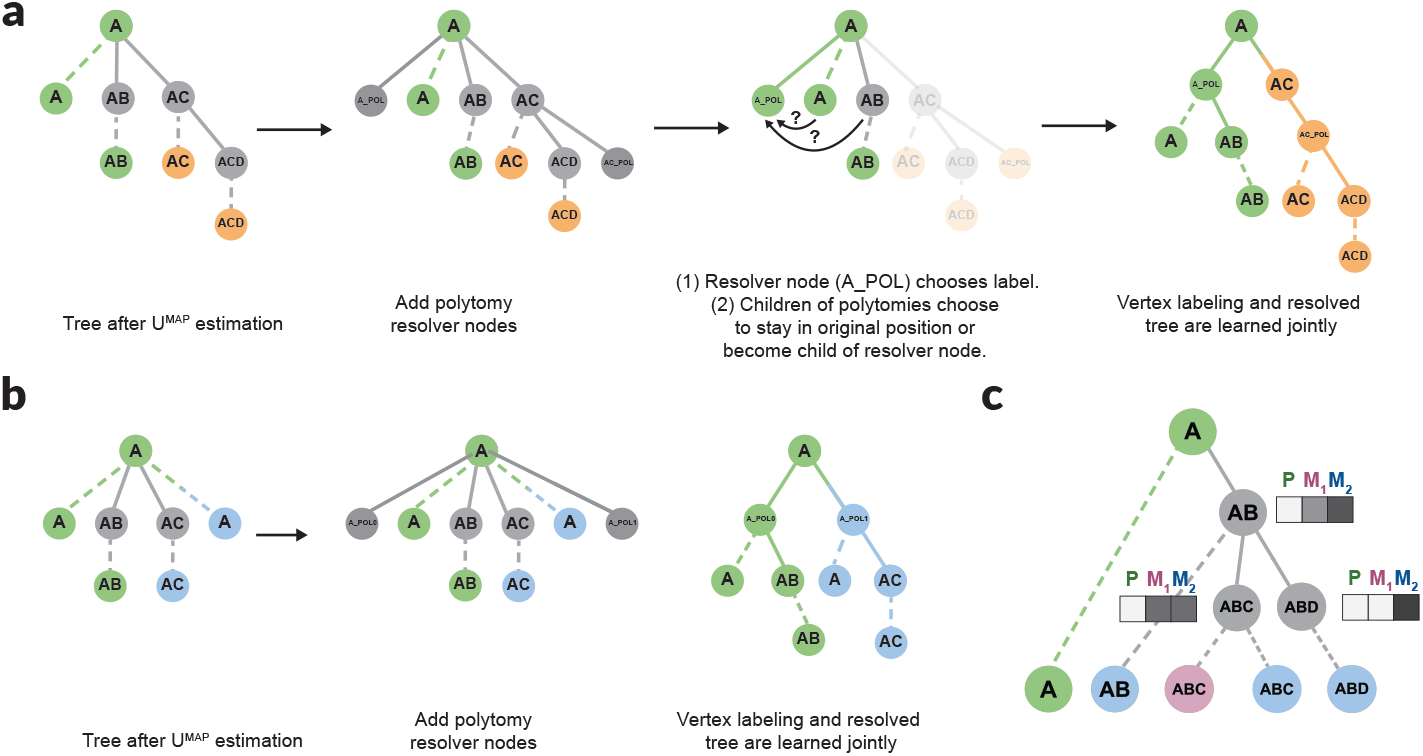
**(a)** Polytomy resolution algorithm with two nodes (A and AC) that have polytomies that can be resolved. **(b)** Polytomy resolution algorithm for a single node with four children and thus two resolver nodes. **(c)** Weight initialization is done such that nodes start with higher probabilities of being in the same site as the site that they or their children are detected in (after **U**^MAP^ estimation).

**Figure S12.**
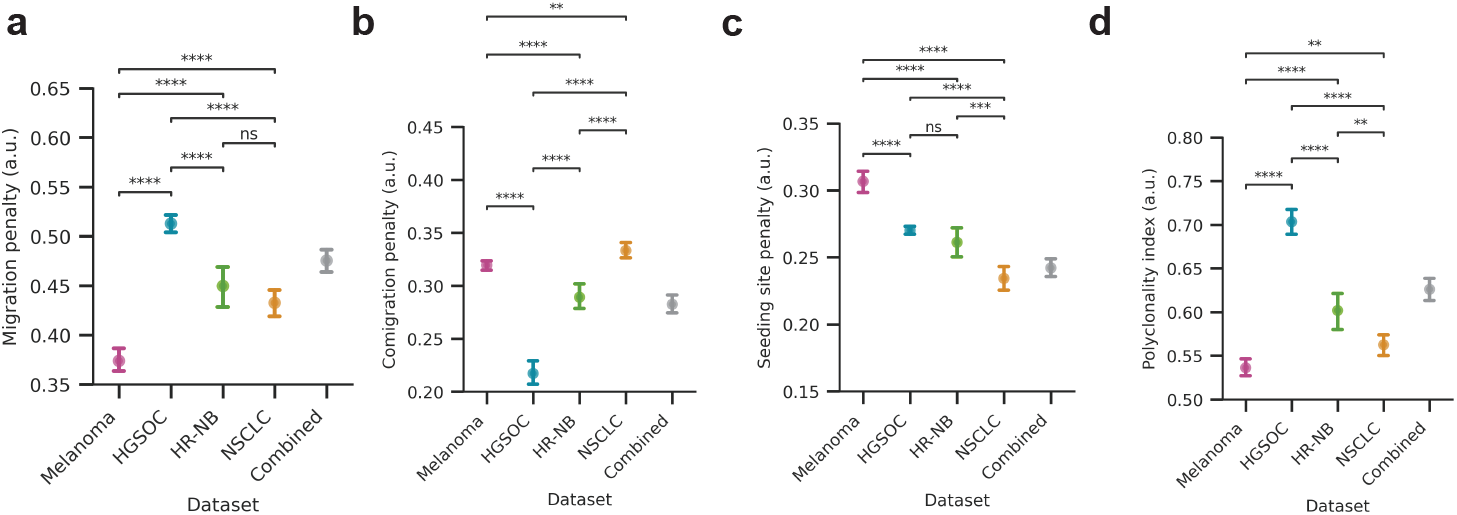
The **(a)** migration penalty/weight, **(b)** comigration penalty/weight, and **(c)** seeding site penalty/weight for each cohort, when taking 100 bootstrap samples of each cohort and fitting the weights to the bootstrapped sample. **(d)** The polyclonality index, which is 1 − (*w*_*c*_/(*w*_*m*_ + *w*_*c*_)), where *w*_*m*_ is the migration penalty/weight and *w*_*c*_ is the comgiration penalty/weight. Statistical significance tested through a Welch’s t-test; ns: 5e-02 < p <= 1, *: 1e-02 < p <= 5e-02, **: 1e-03 < p <= 1e-02, ***: 1e-04 < p <= 1e-03, ****: p <= 1e-04. Error bars are the standard error for each dataset.

**Figure S13.**
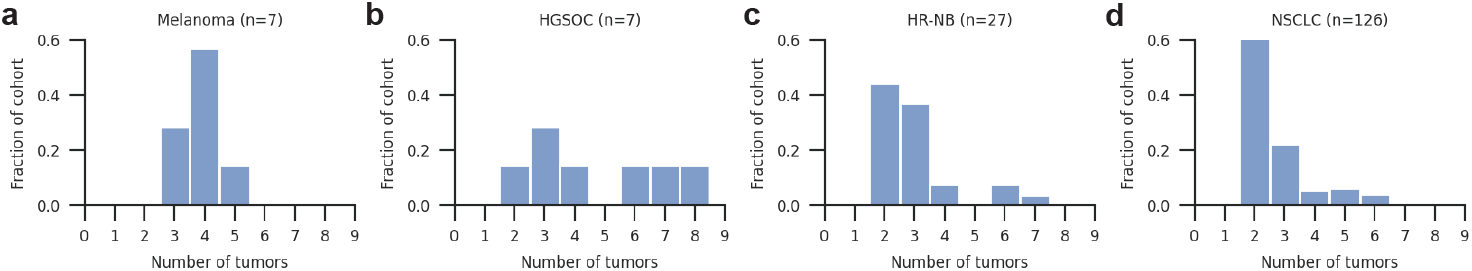
The distribution of tumors (number of distinct anatomical sites) for each cohort: **(a)** melanoma, **(b)** high-grade serous ovarian cancer (HGSOC), **(a)** high-risk neuroblastoma (HR-NB) and **(a)** non-small cell lung cancer (NSCLC).

**Figure S14.**
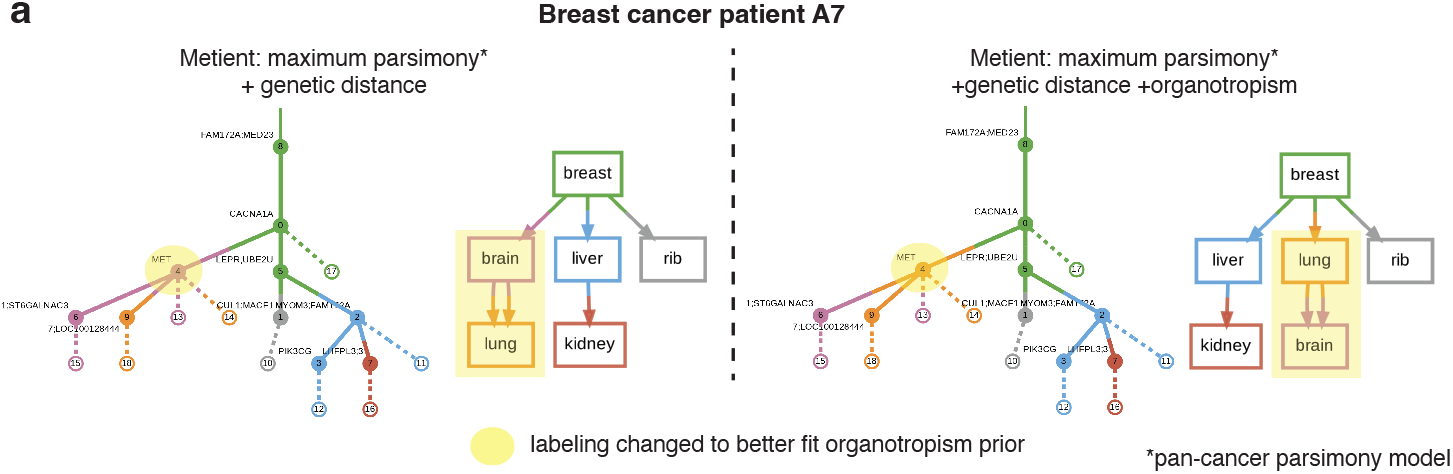
Metient’s organotropism prior corrects unlikely patterns of seeding. **(a)** The inferred migration history for breast cancer patient A7^68^ without (left) and with (right) the inclusion of the organotropism prior. The addition of an organotropism prior changes the vertex labeling of clone 4 from originating in the brain to originating in the lung. Solid edges are edges in the clone tree, and dashed edges indicate the presence of the clone in the corresponding colored anatomical site (i.e., witness nodes).

### A. Evaluating migration histories

We present our technique for optimizing migration histories in the context of variational inference. Our goal is to approximate the conditional density of latent variable **V** given observed variables **U** and **T**: *p*(**V** | **U, T**). **U** has been optimized as described in the section “Estimating observed clone proportions” in Methods. *p*(**V** | **U, T**) can be written as:

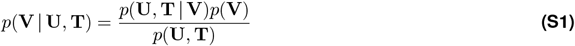

We cannot calculate the denominator, or the evidence, as its derivation is intractable (there are many possible values of **V**):

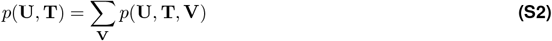

We approximate the posterior distribution *p*(**V** | **U, T**) with a simpler distribution *q*(**V**), and we aim to minimize the Kullback-Leibler (KL) divergence between *q*(**V**) and the true posterior *p*(**V** | **U, T**). The Evidence Lower Bound (ELBO) is given by:

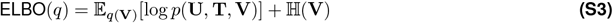

Where the second term is the entropy term.

To handle the categorical nature of **V**, we use the Gumbel-Softmax reparameterization trick to optimize **V**. Starting with a matrix *ψ* ∈ {0, 1}^*K*×*C*^, of randomly initialized values, where *K* is the number of anatomical sites and *C* is the number of clones, and each column represents the unnormalized log probabilities of clone *c* being labeled in site *k*:

1. At every iteration, for each clone *c*, we sample *g*_1*c*_…*g*_*kc*_, *k* i.i.d. samples from Gumbel(0,1) and compute *y*_*ic*_ = *ψ*_*ic*_ + *g*_*ic*_. Where a sample *g* from the Gumbel is computed as:

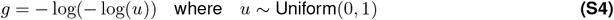
2. We then sample from the categorical distribution represented by the column vector *ψ*_:*c*_ by setting *i*^∗^ = argmax_*i*_ *y*_*ic*_ and represent that sample with a one-hot encoding in **V**, i.e., **V**_*ic*_ =1 if *i* = *i*^∗^, 0 otherwise.
3. Then we evaluate the ELBO(*ν*) where

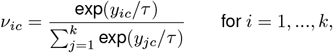

At the end of training, as *τ* approaches 0, then the gradient becomes unbiased and *ν* approaches **V**. In order to capture multiple modes of the posterior distribution, each representing different hypotheses about the migration history, we optimize multiple **V**s in parallel. To do this, we set up steps 1-3 such that *x ψ*s are solved for in parallel ^65^ (with a different random initialization for each parallel process), where *x* is equal to the sample size and is calculated according to the size of the inputs (*α K*^*C*^).

Using the Gumbel-Softmax reparameterization as described above, we approximate the expectation in the ELBO with a sample of **V**, which we denote 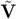:

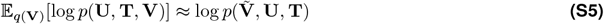

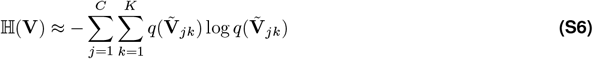

In the following sections, we describe how we calculate 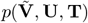, which is broken down into (1) 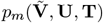, i.e., the scoring of 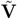 using maximum parsimony, (2) 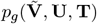, i.e., the scoring of 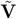 using genetic distance, and (3) 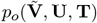, i.e., the scoring of 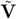 using organotropism.

#### A.1. Evaluating maximum parsimony

As previously described by MACHINA ^17^, the maximum parsimony metrics are defined as:

- **migration number** *m*: Given clone tree **T** and clone tree labeling **V**, the migration number is the number of edges in **T** where the outgoing node and incoming node have a different label. It is the number of edges in migration graph **G**.
- **comigration number** *c*: Given clone tree **T** and clone tree labeling **V**, the comigration number is a subset of the migration edges between two anatomical sites, such that the migration edges occur on distinct branches of the clone tree. It is the number of multi-edges in migration graph **G** if **G** does not contain cycles.
- **seeding site number** *s*: Given a clone tree **T** and clone tree labeling **V**, the seeding site number is the number of unique anatomical sites with an outgoing edge. It is the number of edges in migration graph **G** with an outgoing edge.

Maximum parsimony scoring calculates the number of migrations *m*, comigrations *c*, and seeding sites *s*.

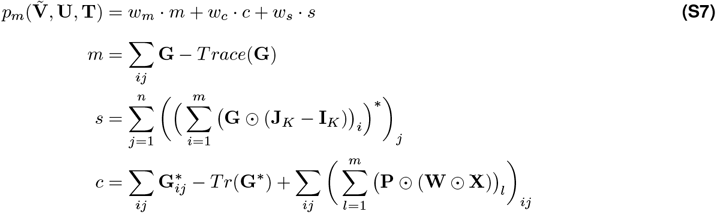

where 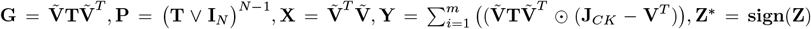. V represents boolean matrix multiplication, **I**_**n**_ is a *n* × *n* identity matrix, ⊙is the Hadamard, i.e., element-wise product, and **J**_*mn*_ is a matrix of ones with dimensions *m* × *n*.

#### A.2. Evaluating genetic distance

Genetic distance is a measure of the number of mutations between clones. Given a distance matrix **D** which has normalized genetic distances between every clone:

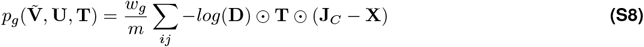

where **J**_*C*_ is a square matrix of ones, ⊙ is the Hadamard, i.e., element-wise product, and 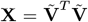. The product **T** ⊙ **J**_*C*_ − **X** tells us if two nodes have an edge between them and they are in different sites. Taking the hadamard product of this with the negative log of **D** gives lower scores to edges with higher genetic distances. We normalize by the migration number *m* so we don’t implicitly penalize migration histories with more migrations through this scoring.

#### A.3. Evaluating organotropism

Organotropism refers to the observation that certain cancers metastasize to specific organs. We penalize migration edges between organs that are less likely to occur based on clinical data. Given a vector **o** which contains the frequency that a primary tumor seeds other anatomical sites:

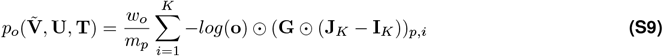

where 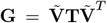, ⊙ is the Hadamard, i.e., element-wise product, **J**_*K*_ is a square matrix of ones, and **I**_**K**_ is the identity matrix. The product (**G** ⊙ (**J**_*K*_ −**I**_*K*_)) contains the number of migrations between different sites, and taking the Hadamard product of this with the negative log of **o** gives lower scores to migration edges with higher organotropism frequencies. The subscript *p, i* represents taking the row of (**G** ⊙ (**J**_*K*_ − **I**_*K*_)) which represents the primary site index and summing over the columns at every other anatomical site *i*. We normalize by *m*_*p*_, the number of migrations originating from the primary site, so we don’t implicitly penalize migration histories with more migrations through this scoring.

### B. Calibrate alignment

A parsimony model is represented by a set of parsimony weights – *w*_*m*_, *w*_*c*_, and *w*_*s*_ – assigned, respectively, to the number of migrations (indicated by *m*), comigrations (*c*), seeding sites (*s*). A migration history’s parsimony score, *p*, is the model-weighted average of these three parsimony metrics, i.e., *p* = *w*_*m*_*m* + *w*_*c*_*c* + *w*_*s*_*s* (Equation S7). Different parsimony models favor different histories on the Pareto front. To fit a parsimony model to a cancer type-specific cohort, we look at how well the maximum parsimony distribution aligns with the genetic distance distribution of each patient’s migration history trees.

Take a cohort of *N* patients, where each patient, *n*, is associated with a set,

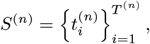

of *T* ^(*n*)^ migration histories. Each migration history *t* is associated with a genetic distance *g*_*t*_ (or, alternatively, an organotropism score), and a vector of parsimony metrics **x**_*t*_ = [*m*_*t*_ *c*_*t*_ *s*_*t*_] (i.e., the counts of migrations, comigrations, and seeding sites, respectively). The goal is to set the parameters, *θ* = [*w*_*m*_ *w*_*c*_ *w*_*s*_] of the parsimony prior 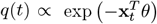 so that it matches, as best as possible, a target distribution, *p*(*t*), over the migration histories *t* implied by the *g*_*t*_, where *p*(*t*) *α* exp(− *τg*_*t*_) and *τ* is a user-defined “temperature” hyper-parameter.

To fit these parameters, we define patient-specific categorical distributions *p*^(*n*)^(*t*) and *q*^(*n*)^(*t*) as follows. Let **g**^(*n*)^ be the vector of length *T* ^(*n*)^ of genetic distances of the migration histories for patient *n*, where 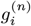 is the genetic distance for the *i*-th tree. And let the column vector 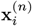 be the parsimony metrics for the *i*-th migration history associated with patient *n*. We will append the *T* ^(*n*)^ vectors 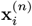 to make a 3 × *T* ^(*n*)^ design matrix *X*^(*n*)^. Also we define the vector-valued softmax function in the typical way, i.e.,

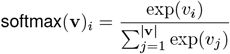

where softmax(**v**)_*i*_ is the *i*-th element of the vector output by softmax(**v**). Then the “parsimony” probability distribution over the trees for patient *n* is represented by the vector **q**^(*n*)^

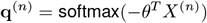

and the target distribution by the vector **p**^(*n*)^

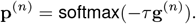

Then we define the cohort calibration objective *E*(*θ*) as an average cross-entropy over the patient cohort, i.e.,

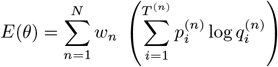

and the MLE estimate of the parameters is *θ*^∗^ = argmax_*θ*_*E*(*θ*). *w*_*n*_ is set to *log*(*E/*(*r* · *b*)), where *E* is the number of internal edges of a patient’s clone tree, *r* is the number of possible primaries for the patient, and *b* is the number of possible clone trees for a given patient (so as not to bias towards patients with multiple possible primaries or multiple possible clone trees). Since the number of edges is equal to the maximum number of migrations possible in a tree, it is also equal to the number of possible genetic distance observations that that tree can provide in the genetic distance scoring of that tree. Therefore, *w*_*n*_ is representative of the information content that a patient can provide when fitting *θ*.

#### B.1. Specifying the target distribution by setting the temperature parameter

The use of *E*(*θ*) to set *θ* requires that for a patient *n* that, generally speaking, the genetic distance 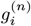 for a potential migration history, represented by a tree *i*, is lower for more probable histories. However, because *E*(*θ*) is minimized when *τ* **g**^(*n*)^ = *θX*^(*n*)^ + *c***1** for some constant *c*, this could be a very strong assumption, one that we might not always be comfortable making.

Fortunately, we can set *τ*to increase the correctness of this assumption. Notice that in the limit of large *τ* that

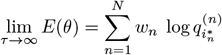

where 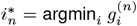, assuming that the minimum is unique. If the minimum is not unique then the above is true if we replace 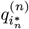 with the average of 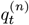 of all the trees *t* that have the minimum genetic distance for patient *n*.

So, in other words, if we set *τ* to be very large, then *E*(*θ*) is just the (weighted) sum of the log probabilities of the minimum genetic distance trees in each patient, and optimizing *E*(*θ*) corresponds to maximizing the parsimony probabilities of the best scoring trees per patient under the genetic distance score.

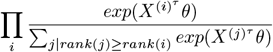

So, we set *τ* to be large, such that *τ* is multiple times the maximum genetic distance (assuming that the genetic distance is always positive). We do the same for the organotropism prior.

### C. Case-by-case differences to expert annotations

#### C.1. Comparisons to Melanoma patients from Sanborn et al

Migration histories generated for the metastatic melanoma cohort using Metient-calibrate agree with the expert analysis that most melanoma patients exhibit primary single-source seeding (Supplementary Figure S7). For patient F (Supplementary Figure S7c), our reconstruction of the clone tree and observed clones does not suggest that a lymph node to distant metastasis seeding event is likely, but that this patient also likely exhibits a primary-only seeding pattern. In the best solution predicted for patient D, Metient predicts that a locoregional skin metastasis from the right ankle could have given rise to subsequent metastases, supporting one of the possible paths (in dotted lines) that the original authors propose (Supplementary Figure S7d). The second best solution shows primary-only seeding, which is another possible path proposed by the authors (Supplementary Figure S7d).

#### C.2. Comparisons to HGSOC patients from McPherson et al

In the seven HGSOC patients, predicted migration histories by McPherson et al. ^4^ were made available using an algorithm that only minimizes migrations (Sankoff algorithm ^19^). We find that four out of seven patients are in complete agreement (Supplemental Figure S8). For patient 1, by resolving polytomies, we offer an explanation with less migrations and comigrations, and predict that the left fallopian tube rather than the small bowel served as a possible intermediate site before further metastatic dissemination (Supplemental Figure S8a). For patient 3, we offer an explanation with less migrations, comigrations and seeding sites, suggesting that all metastases were seeded from the primary (Supplemental Figure S8c). Finally for patient 7, solving for polytomies allows us to reduce the migration number by 1 from the right uterosacral to left ovary, although the overall seeding pattern is in agreement (Supplemental Figure S8d).

#### C.3. Comparisons to HR-NB patients from Gundem et al

Because the HR-NB annotations only indicate the presence of a migration between two sites and not the directionality, we compared our site-to-site migrations (i.e., a binarized representation of migration graph **G** (Figure S1c)) to those that were previously reported. We looked at the 14 HR-NB patients for which there were manual expert annotations from Gundem et al. ^9^, and found that we predict the same overall site-to-site migrations for 10 out of 14 cases. For patient H103207, we predict their before therapy pattern on the Pareto front (Solution 3 in Figure S5a), but we prioritize two solutions with metastasis-to-metastasis seeding between the lung and the liver. A subset of this seeding between the liver and two lobes of the lung is suggested in their after therapy hypothesis of spread (Figure S5a). While Gundem et al. suggest seeding between the two lobes of the lung as well as from each lobe of the lung to the liver, we infer a simpler, serial progression, where the right lung lower lobe seeds the liver, which subsequently seed the left lung lower lobe (Solution 1 in Figure S5a). For patient H132396, Metient prioritizes migration histories with fewer migrations (Solutions 1 and 2 in Figure S5g), but presents the expert annotation on the Pareto front (Solution 3 in Figure S5g). For patient H132384, Metient proposes bone-to-bone secondary metastasis formation (Solution 1 in Figure S6d), but again presents the expert annotation on the Pareto front (Solution 2 in Figure S6d). For patient H134821, we infer the same pancreas to hilar lymph node seeding proposed by the authors as spread after therapy, as well as the same seeding from the primary to the cervical metastasis. However, we propose that rather than the primary tumor seeding both the cervical and thoracic lymph nodes, the primary tumor likely seeded either the thoracic lymph node (Solution 1 in Figure S6f) or the cervical lymph node (Solution 2 in Figure S6f) which then leads to further dissemination.

### D. Model choice impacts downstream analyses

As we were analyzing different aspects of metastatic dissemination, we asked how these answers might change if a seeding model is enforced when reconstructing a patient’s migration history. To highlight how the choice of seeding model can impact the analysis and interpretation of metastatic dissemination, we compared the migration histories produced by three models: (1) assumption of primary, single-source seeding, (2) the MACHINA assumptions, which first minimize migrations, and then break ties based on comigration number followed by seeding site number, and finally (3) the adaptive Metient model fit to each cohort. As expected, a primary, single-source seeding model chooses a primary, single-source dissemination pattern for 100% of patients (Supplementary Figure S4d). The migration penalizing model chooses a primary single-source seeding explanation in 82.6% of patients, and Metient falls in between the two, choosing a primary single-source seeding explanation in 86.2% of patients (Supplementary Figure S4e). Importantly, since Metient can recover and evaluate the relative trade-offs of the parsimony metrics, when choosing a primary single-source solution, our model has either not found a plausible metastasis-to-metastasis explanation for a patient’s data on the Pareto front, or has used the metastasis priors to deem such an explanation less likely. In contrast, previous models do not automatically recover multiple possible hypotheses, therefore reducing confidence in these algorithms’ choice of best history.

In addition to having an impact on the inferred seeding patterns, a model that assumes primary single-source seeding also changes other interpretations of metastatic seeding. We asked two questions about the best migration histories produced by the two extremes of models, i.e. the assumption of primary, single-source seeding and Metient: (1) the frequency in which a new seeding site is added, and (2) the frequency of polyclonal migrations between two sites. As expected, a model which assumes primary, single-source seeding promotes migration histories with only one seeding site (Supplementary Figure S4f). In turn, such a model infers a higher fraction of polyclonal migrations (Supplementary Figure S4b) compared to the histories prioritized by Metient. The trade-off between polyclonality and seeding sites occurs because additional seeding sites reduce the number of migration edges that must be placed between the primary and all other metastases. Balancing this trade-off correctly is important as it impacts the interpretation of seeding clonality as well as which clones perform seeding. Specifically, 9% (15/167) of patients have differing colonizing clones between the two models, changing the inference of which clones, and therefore which mutations, have metastatic competence.

### E. Pre-processing of sequencing data

#### E.1. Variant read probability calculation (ω)

In order to account for non-diploid copy number and tumor purities, we require a variant read probability *ω* to be input for every genomic locus in each sample. For a given sample *s* and variant allele *j*, the variant read probability *ω*_*js*_ is the probability of observing a read with the variant allele at that locus in a cell with the mutation, and is calculated as:

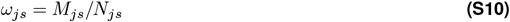

where *M*_*js*_ is the number of copies of the mutant allele *j* in sample *s* in the cells that contain the mutant allele, and *N*_*js*_ is the average number of copies at the genomic locus of the mutation *j* in all cells in *s*.

To account for the fact that cancer cells frequently have different numbers of copies at genomic loci compared to normal cells, *N*_*js*_ is calculated as:

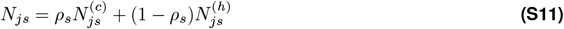

where:

- 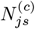 is the population average copy number of the locus which contains mutant allele *j* in the cancer cell population
- 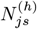 is the copy number at the genomic locus of mutation *j* in the normal cell population. In diploid cells this is 2, and in haploid cells this is 1.
- *ρ*_*s*_ is the tumor purity of sample *s*

*ρ*_*s*_ and 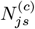 (and sometimes *N*_*js*_) are normal outputs from a copy number calling pipeline. We suggest setting *M*_*js*_ =1 unless there is strong evidence that the *j* allele has been amplified. In this case, allele-specific copy number callers provide the major allele copy number *A*_*js*_ and minor allele copy number *B*_*js*_, where 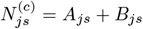, and *M*_*js*_ = *A*_*js*_. When a locus is impacted by many different CNAs, accurately estimating *M*_*js*_ is challenging since there are likely subclonal changes in the multiplicity of the *j* allele, in which case we recommend excluding these mutations. For additional information on how to estimate *M*_*js*_ and *N*_*js*_ please refer to Tarabichi et al. ^69^.

If clustering is used, we have to properly combine multiple SNV loci with different potential variant read probabilites. To do this, we rescale the reference and variant allele read counts for each locus and then set its variant read probability to 0.5 before combining variants within a cluster (where we add the reference and variant allele read counts for all variants within a cluster). This rescaling allows us to effectively treat the variant as coming from a diploid locus. To achieve this, we use the following rescaling formulas, which has been previously described in Wintersinger et al. ^58^:

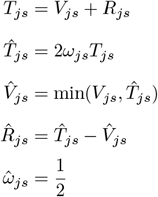

Where *T*_*js*_ is the input count of total reads, *V*_*js*_ is the input count of variant reads, *R*_*js*_ is the input count of reference reads, and *ω*_*js*_ is the variant read probability at a genomic locus *j* in anatomical site *s*. The rescaled total, reference, and variant allele read counts and variant read probability are 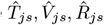 and 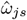 respectively.

#### E.2. Breast Cancer Dataset

The single nucleotide variant calls from two breast cancer patients with whole genome sequencing data were taken from Hoadley et al. ^68^. The variant calls were in copy number neutral variant positions and tumor purity was not reported, so reference and variant counts along with defaults for tumor purity, major copy number and minor copy number (defaults are 1.0, 1, 1, respectively) were inputted into PyClone-0.13.1 clonal analysis ^70^. PyClone’s MCMC chain was run for 100,000 iterations, discarding the first 50,000 as burnin. Orchard was run using the PyClone clusters as input with -p flag to force trees to be monoprimary (come from a singular root cancer clone) and all variant read probabilities set to the default of 0.5, since SNVs from regions with CNAs were excluded, and tumor purity was not reported and thus assumed to be 1. We ran Metient-evaluate on this data using all default configurations (dynamically calculated sample size based on size of input clone tree and number of anatomical sites).

#### E.3. High-grade Serous Ovarian Cancer Dataset

To better compare to McPherson et al.’s own migration history analysis, we used the mutation clusters, clone trees and cellular prevalences of each clone that they estimate and report ^4^. Metient was run with the **U** matrix inputted, and we solve for **V** for each patient. We ran Metient-calibrate on this data using all default configurations (dynamically calculated sample size based on size of input clone tree and number of anatomical sites) and with polytomy resolution.

#### E.4. Melanoma Dataset

The single nucleotide variant and copy number calls from eight melanoma patients with whole exome sequencing data were taken from Sanborn et al. ^3^, along with estimated tumor purity. Only SNVs in copy number neutral regions were considered. Patient H was excluded due to a lack of copy number neutral SNVs. Reference and variant read counts along with major and minor copy number and tumor purity were inputted into PyClone-VI 0.1.3 for clonal analysis ^71^. PyClone-VI’s fit command was run with all default parameters. Orchard was run using the PyClone clusters as input with -p flag to force trees to be monoprimary (come from a singular root cancer clone). Variant read probabilities for Orchard were calculated using major copy number, minor copy number and tumor purity according to Equation S10. We ran Metient-calibrate with the clonal proportions estimated by running Orchard (i.e., *÷* in Orchard’s output) using all default configurations and with polytomy resolution.

#### E.5. Neuroblastoma Dataset

Access to multi-WGS data for 45 neuroblastoma patients was provided through dbGaP accession phs03111^9^. Of these 45 patients, 27 patients had at least one primary and one metastatic tumor sample with a tumor purity of >10%, and all analysis was conducted on this patient subset. Single nucleotide variant, copy number calls and tumor purities were collected from this dataset, and clusters produced from the original paper using DPClust ^72^ were used. Multiple samples for the same anatomical site and sample time (i.e., diagnosis, therapy-naive re-resection, therapy resection during induction chemotherapy, relapse or further relapse) were combined by pooling reference and variant allele counts. Orchard was run using the DPClust clusters as input with -p flag to force trees to be monoprimary (come from a singular root cancer clone). Variant read probabilities for Orchard and Metient were calculated using major copy number, minor copy number and tumor purity according to Equation S10. We ran Metient-calibrate with the clonal proportions estimated by running Orchard (i.e., *η* in Orchard’s output) using all default configurations and with polytomy resolution.

For three patients (H103207, H132388, H134822), multiple primary tumor samples were collected at different time points (diagnosis and resection during therapy). For these patients, we treated the therapy resection and diagnosis tumor as multiple samples from the same anatomical site if the anatomical site was labeled the same, and as two different primaries if the anatomical sites were different. The therapy resections were usually taken a few months after diagnosis tumor samples.

#### E.6. Non-small Cell Lung Cancer Dataset

We used the clustered SNVs, clone trees and observed clone proportions made available by the TRACERx consortium for 126 non-small cell lung cancer (NSCLC) patients (downloaded from https://zenodo.org/record/7649257). When samples for multiple regions of a tumor were available, the reference and variant allele counts were summed together to generate reference and variant allele counts for the entire tumor. Since we model variant allele counts as binomially distributed with *n* total reads (variant + reference) and *p* probability of generating a variant read, this summing assumes that each sampled region of a tumor has the same probability *p*. Metient was run with the **U** matrix inputted, and we solve for **V** for each patient. We ran Metient-calibrate on this data using all default configurations (dynamically calculated sample size based on size of input clone tree and number of anatomical sites) and with polytomy resolution.

#### E.7. Single-cell lineage tracing data

Single-cell lineage tracing data from a Cas9-based lineage recorder was taken from Quinn et al. ^22^. We pre-processed the barcode and scRNA-seq data from the 100 implanted tumor clones using Cassiopeia ^73^, using the Vanilla Greedy Solver to generate trees for each clone. We ran Metient and FitchCount ^22^ on these trees, where the presence of each node in the tree was taken directly from the sequencing data. Metient was run in evaluate mode with a sample size of 38,000 to account for the large input sizes, using the parsimony model weights that were calibrated to the collective bulk DNA sequencing cohort of 167 patients from the melanoma, HGSOC, HR-NB and NSCLC cohorts. Organotropism frequencies were also inputted, by computing the number of times each of the 100 clones metastasized to a tissue.

For the gene expression analysis of clones 95 and 97 in Figure 6d and Figure 6f, scanpy ^74^ was used. The gene by cell matrix for each clone was pre-processed by removing genes expressed in less than three cells, normalizing by the median UMI count, and log-transforming the data. Cosine similarity of gene expression was compared between the cell of interest and all other cells in the same clone, in each tissue. To determine the tissue of origin for all cells, we used the top migration history from Metient, and the randomly chosen migration history returned by FitchCount.

### F. Validation of organotropism prior in metastatic breast cancer cohort

To validate the organotropism prior, we ran Metient, using the pan-cancer parsimony model, on samples available from two patients with metastatic breast cancer ^68^ where site labels could be mapped to those used in our organotropism matrix. When faced with multiple parsimonious migration histories, Metient chooses a more plausible tree, wherein lung to brain seeding is preferred over brain to lung seeding, which is clinically rare ^41^ (Figure S14a).

